# Stimulus-Driven Thermodynamic Shifts and Geometric Reorganization in Mouse Primary Visual Cortex

**DOI:** 10.64898/2026.02.12.705646

**Authors:** Jirui Liu, Longsheng Jiang, Xueting Li, Jiamin Wu, Jia Liu

**Author notes:** Correspondence: J.L.

## Abstract

Efficient coding is essential for sensory systems to extract meaningful information from the environment. Here, we investigate how stimulus-driven thermodynamic shifts and geometric reorganization enable efficient population coding. Using wide-field calcium imaging, we simultaneously recorded neuronal activity across the entire mouse V1 under the presentation of structured stimuli and characterized neural dynamics spanning microscale neuronal connectivity, mesoscale thermodynamic states, and macroscale manifold geometry. We found that stimulus presentation reorganized neuronal couplings into increasingly modular subnetworks, driving a shift from near-critical dynamics toward a more ordered regime. This shift coincided with a compression of neural population activity onto low-dimensional manifolds aligned with stimulus features, thereby enabling efficient coding. Furthermore, mathematical derivations and *in silico* perturbation experiments confirmed that selective modulations of neuronal connectivity altered both critical temperature and manifold geometry, establishing a causal link from neuronal couplings to efficient coding. Collectively, our findings suggest a mechanistic bridge between statistical physics and neural geometry, providing new theoretical insights into how neural networks dynamically transition into more stable and efficient coding states in response to the environment.

## Introduction

Natural sensory inputs are richly structured and highly redundant, yet neural systems must represent them using noisy spikes under stringent metabolic constraints. Efficient coding theory frames this challenge as a normative compression problem: asking how neural populations optimize information transformation per unit of metabolic cost (Atick & Redlich, 1990; Chalk et al., 2018; Laughlin, 2001; Levy & Baxter, 1996; Pitkow & Meister, 2012). Classical theories propose that early sensory processing reduces redundancy by adapting to the statistical structure of natural signals through transformations such as decorrelation and sparse coding, producing compact representations that downstream neural circuits can efficiently decode (Attneave, 1954; Benjamin et al., 2022; Olshausen & Field, 1996). More recent frameworks emphasize efficiency as a dynamic rather than static objective, arguing that optimal coding continuously adapts to changes in stimulus statistics and task demands while balancing information content, robustness, and energetic cost over time (Cai et al., 2025; Młynarski & Hermundstad, 2021; Panzeri et al., 2022).

Modern population-level recordings conceptualize these principles geometrically. Each stimulus or stimulus category evokes a distinct cloud of neural activity in a high-dimensional state space. Collectively, these clouds form a structured neural manifold rather than occupying state space indiscriminately (Stringer et al., 2019), whose dimensionality and geometric organization determine how effectively downstream circuits can discriminate relevant stimulus categories (Chung & Abbott, 2021; Cunningham & Yu, 2014; Langdon et al., 2023; Ma et al., 2025). Empirical and theoretical analyses have consistently shown that untangling of neural manifolds significantly increases linear separability and classification performance (Chung & Abbott, 2021; Cohen et al., 2020; DiCarlo & Cox, 2007; Ma et al., 2025; Rigotti et al., 2013). Critically, neural manifold geometry is shaped by correlated variability among neurons, governed by covariance structures emerged collectively from local neuronal interactions (Churchland et al., 2010; Ponce-Alvarez et al., 2023; Safavi et al., 2024). Sensory inputs rapidly modify these variability structures, reducing trial-to-trial fluctuations and selectively modulating cortical noise correlations (Churchland et al., 2010; Ecker et al., 2010; Renart et al., 2010). Consequently, a key unresolved question remains: how do microscale neuronal interactions shape macroscale neural manifold geometry during sensory processing?

Thermodynamics provides a principled framework for understanding how changes in interaction structures affect global fluctuations, which subsequently shape the dimensionality and geometry of neural manifolds. Indeed, microscale neuronal interactions, such as effective couplings and shared variability, represent precisely those quantities that statistical physics aims to coarse-grain into macroscale descriptors of collective network states. A prominent illustration of this approach is the criticality framework, inspired by observations of cortical activity exhibiting scale-free cascades (i.e., neuronal avalanches), indicative of operation near a critical phase transition (Beggs, 2008; Beggs & Plenz, 2003; Hengen & Shew, 2025; Liu et al., 2025; Ponce-Alvarez et al., 2023; Schneidman et al., 2006; Shew et al., 2009; Tkačik et al., 2015; Zhang et al., 2025), with some evidence suggesting a slight deviation (Fosque et al., 2021; Liu et al., 2025; Moretti & Muñoz, 2013).

Additionally, large-scale computational models suggest that optimal neural network organization emerges precisely near the boundary between ordered and disordered dynamical regimes (Haimovici et al., 2013; Shew et al., 2015; Zhang et al., 2025).

Specifically, within equilibrium maximum entropy (Ising) models of neural populations, criticality manifests as a distinct peak in specific heat, and the difference between the critical temperature *T_c_* and the network’s operational (working) temperature concisely indicates how closely the fitted interactions place the network near the transition point (Schneidman et al., 2006; Tkačik et al., 2015). Conceptually, a shift in *T_c_* implies alterations in the global fluctuation landscape, specifically influencing how readily neural populations explore various collective states. Thus, this shift directly determines whether neural activity spans many degrees of freedom or becomes confined to a small number of dominant modes, thereby defining the dimensionality of the neural representational space. Recent computational studies underscore the relevance of this thermodynamic perspective for efficient coding, showing that optimal coding performance occurs precisely at noise levels coinciding with critical-like signatures (Safavi et al., 2024). Accordingly, we propose that sensory inputs restructure microscale neuronal interactions along a thermodynamic continuum, resulting in a redistribution of population variance into a lower-dimensional manifold aligned with stimulus features, thus increasing efficient coding.

To test this hypothesis, we employed a multi-scale approach connecting microscale neuronal interactions, mesoscale network dynamics, and macroscale coding geometry. At the microscale, we utilized RUSH3D, a recently developed high-speed 3D mesoscopic imaging system (Zhang et al., 2024), to simultaneously record calcium activity from thousands of neurons in mouse primary visual cortex (V1).

RUSH3D provides a centimeter-scale volumetric imaging with single-neuron resolution and minimal phototoxicity, enabling continuous *in vivo* recordings for hours. This allowed us to monitor fine-grained neuronal activity patterns during visual stimulus presentation, thus quantifying pairwise neuronal couplings fundamental to network dynamics. At the mesoscale, we fitted pairwise maximum entropy (MaxEnt) models to the recorded population activity to determine how sensory inputs shift the network’s thermodynamic state. Pairwise MaxEnt models define probability distributions over neural activity patterns by maximizing entropy under constraints matching observed mean firing rates and pairwise correlations (Jaynes, 1957; Roudi et al., 2009; Schneidman et al., 2006; Shlens et al., 2006; Tkačik et al., 2010). This approach allowed us to infer effective coupling parameters and assessed whether the network operated near a critical state or transitioned toward a more ordered regime during stimulus presentations. To visualize these thermodynamic transitions, we constructed functional connectivity graphs based on correlations and quantified changes in community structures, testing whether sensory inputs reorganize mesoscale network architectures into modular neuronal clusters. At the macroscale, we explored the geometry of neural population coding by evaluating the effective dimensionality of neural state space and employing support vector machine (SVM) classifiers to quantify stimulus discriminability. Increased decoding accuracy paired with reduced dimensionality would indicate efficient coding, signifying that the network reliably represents stimuli using fewer dimensions. Finally, having established a link between thermodynamic state changes and improved coding efficiency, we tested causality through mathematical derivations and *in silico* perturbation experiments. Overall, this integrative approach validates our hypothesis, demonstrating that sensory inputs drive neural circuits into a more ordered regime, thus facilitating efficient coding.

## Results

### Efficient coding at macroscale level in mouse V1

To probe efficient population coding under structured visual inputs, we employed a 2×2 factorial stimulus design based on shape features (Fig. 1A). Petal-like shapes were defined by two orthogonal features: petal length (deep versus shallow, D&S) and petal width (narrow versus wide, N&W), thereby forming a 2D feature space (Frank et al., 2023). This design yields four discrete, non-overlapping shape categories, each comprising 10 exemplars. In this shape space, categories along either length or width axis were linearly separable. In contrast, diagonal groupings, spiky (deep and narrow, D&N) paired with stubby (shallow and wide, S&W) versus the opposite diagonal pair (deep and wide, D & W; shallow and narrow, S&N), represented a classic XOR problem. Specifically, the XOR classification required distinguishing categories situated at opposite corners along diagonals, which cannot be separated by a single linear boundary in the 2D feature space.

**Figure 1.**
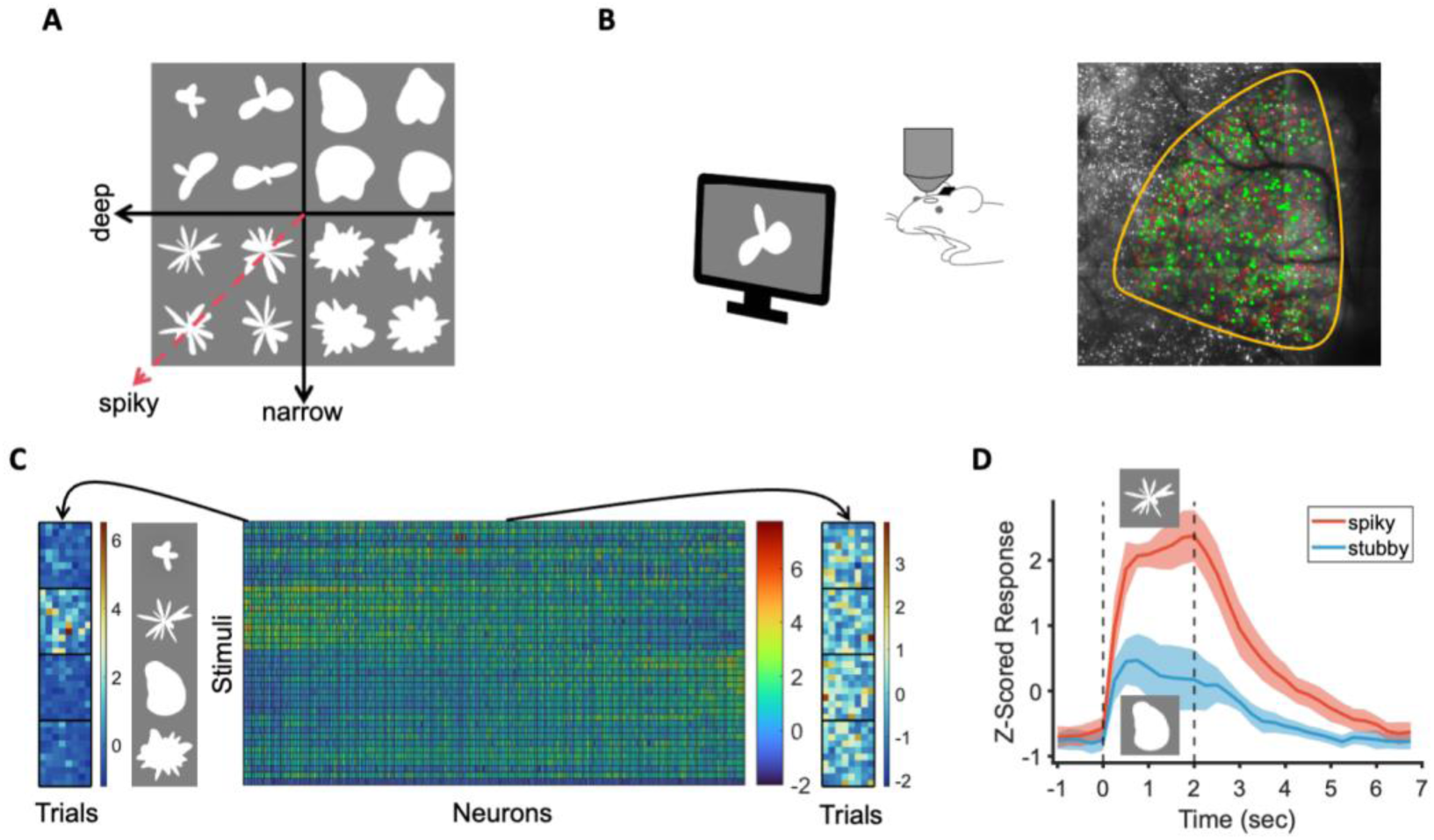
Neuronal and population-level responses in mouse V1. (A) Schematic of the 2D stimulus space defined by shape features (deep/shallow × narrow/wide). This space contains 4 discrete shape categories, each comprising 10 distinct shapes sharing identical feature parameters but with inherent variability. Middle: Mice passively viewed randomly ordered stimulus sequences, where each stimulus was presented for 2 s followed by a 6 s interstimulus interval (ISI) displaying a gray screen. (B) Illustration of wide-field calcium imaging from mouse V1 (region outlined in yellow). Colored circles represent recorded neurons; green circles indicate responsive neurons, and green dots specifically mark R&R neurons. For visual clarity, only a subset of neurons is shown. (C) Middle: Time-averaged responses of all R&R neurons from an example mouse to each of the 40 individual stimuli (rows) during the stim-on epoch. Each column corresponds to one neuron, averaged across eight trials; Left and right panels: Single-trial peak responses at stimulus offset for an example neuron selective to spiky shapes (left) and a non-selective neuron (right). Each matrix represents 40 stimuli (rows) over eight trials. Color intensity indicates the magnitude of peak neuronal responses. (D) Average temporal response profile of all spiky-selective neurons from an example mouse. Red represents responses to preferred stimuli; blue indicates nonpreferred stimuli. Shaded regions denote standard deviation (SD) across neurons; Dashed vertical lines mark stimulus duration.

Wide-field calcium imaging of awake, head-fixed mice revealed robust, stimulus-locked activity throughout V1 (Fig. 1B). Across all recorded V1 neurons (N = 8046 ± 2131 per mouse, 10 mice total), we restricted subsequent analyses to neurons that exhibited responsiveness to at least one stimulus and consistent response patterns across trials, hereafter referred to as reliably responsive (R&R) neurons (Marshel et al., 2011) (N = 199±47 per mouse; defined using criteria for responsiveness and trial consistency detailed in Methods). On average, calcium fluorescence in R&R neurons during the stimulus presentation (stim-on epoch, 0-3.5 s post-stimulus onset; Methods) was elevated by 14±4% relative to the spontaneous epoch (stim-off epoch, 3.5-8 s). This stimulus-driven enhancement is consistent with prior electrophysiological, two-photon imaging, and intrinsic optical signal studies, reinforcing previously characterized visually evoked responses in mouse V1 (Billeh et al., 2020; Bolaños et al., 2024; Carandini et al., 1997; Grinvald et al., 1986; Marshel et al., 2011; Movshon et al., 1978).

We quantified neuronal responses using z-scored activity within the stim-on epoch (Methods) and found that neurons exhibited varied response preferences across different stimulus categories (Fig. 1C, middle). While some neurons did not demonstrate clear selectivity (Fig. 1C, right), others showed distinct categorical tuning (Fig. 1C, left). Additionally, certain neurons specifically encoded individual stimulus features. For example, in the diagonal comparison between spiky and stubby shape categories, a subset of neurons (7.98±3.22% of R&R neurons) showed pronounced selectivity for spiky shapes (Fig. 1D).

This pattern became more evident at the population level, where neural responses to distinct stimulus categories formed separable point clouds in neural state space. Projecting these response clouds onto two orthogonal feature axes (deep–shallow and narrow–wide) revealed a neural geometry closely aligned with the stimulus geometry defined by the four stimulus categories. Tracking the temporal evolution of neural geometries demonstrated that the distance between point-cloud centroids substantially increased following stimulus onset and subsequently decreased after stimulus offset, yet the overall neural geometric structure remained consistent throughout (Fig. 2A).

**Figure 2.**
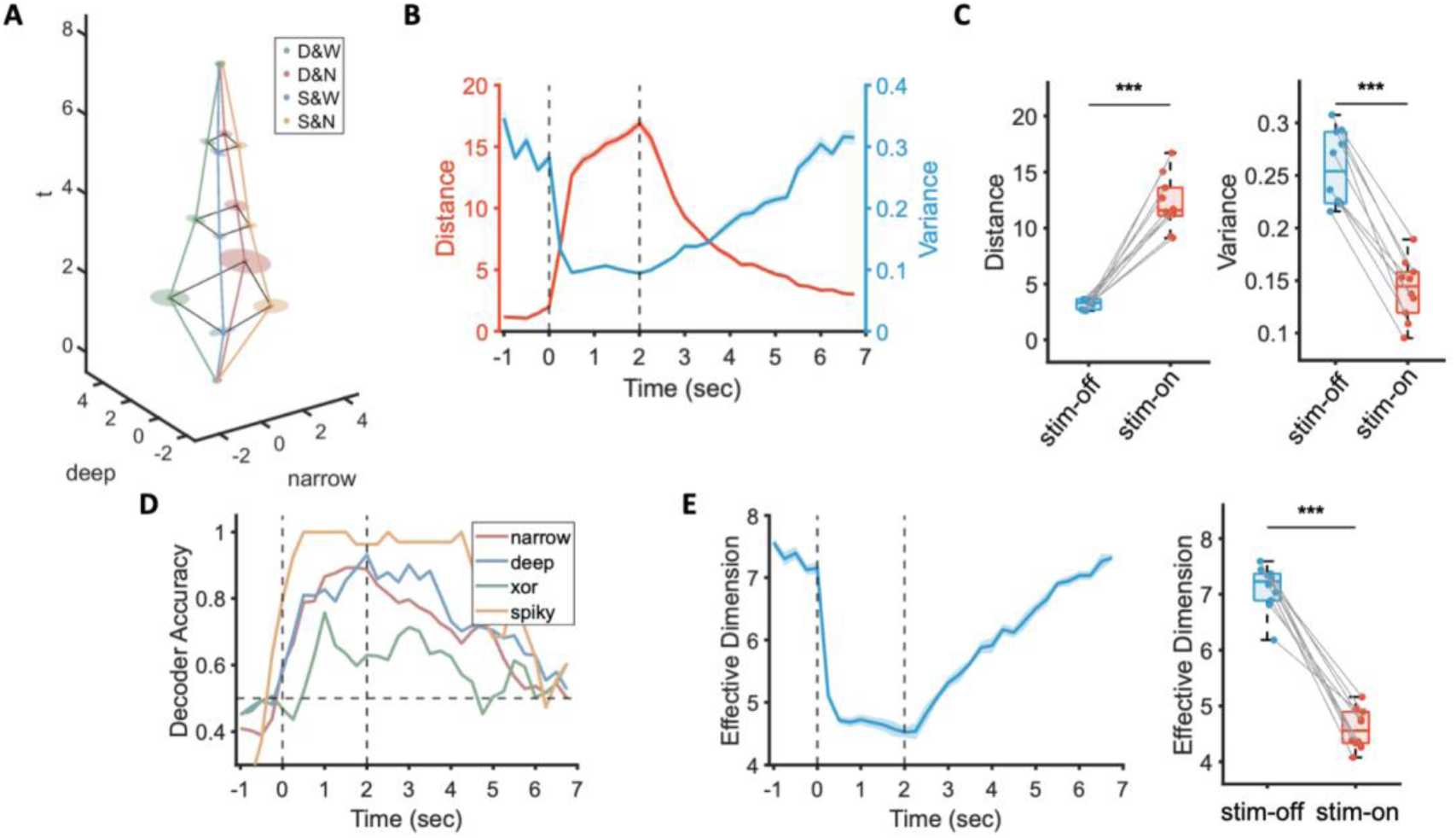
Efficient coding of structured stimulus categories. (A) Temporal evolution of neural population responses to four stimulus categories averaged over trials in an example mouse. Abbreviations: D, deep; S, shallow; W, wide; N, narrow. Point clouds represent neural responses to the 10 stimuli within each category at time points 0, 2, 4, 6 and 8 s relative to stimulus onset, projected onto the two stimulus-feature axes. Shaded regions indicate ±SD across the 10 stimuli within each category. (B) Temporal profiles of inter-cluster distances and intra-cluster variance between spiky and stubby stimulus categories of an example mouse. (C) Inter-cluster distance (Left) and intra-cluster variance (Right) between spiky and stubby categories across 10 mice, measured immediately before stimulus onset (stim-off) and immediately before stimulus offset (stim-on). Values represented averages across all stimuli and trials for each mouse. (D) Temporal dynamics of SVM classification accuracy for pairwise distinctions along four category comparisons in an example mouse. Each feature axis indicates pairwise classifications along that dimension, except for ‘spiky,’ which specifically represents classification of the spiky versus stubby stimuli. Notably, successful categorization in the XOR condition emerges with a substantial delay (Ma et al., 2025). Dashed horizontal line denotes baseline (chance) accuracy level (50%). (E) Left: Temporal dynamics of the effective dimensionality of neural manifolds for an example mouse. Shaded region indicates ±SD from 10 neuron-sampling shuffles. Dashed vertical lines mark stimulus duration. Right: Effective dimensionality of neural manifolds measured immediately before stimulus onset (stim-off) and immediately before stimulus offset (stim-on). Values represent averages across all stimuli and trials from 10 mice. ***: *p* < 0.001.

To quantitatively characterize manifold geometry over time, we used to two point-cloud statistics: centroid distance to measure manifold expansion and contraction and within-cloud dispersion to assess manifold compactness (Fig. 2B). After stimulus onset, the Euclidean distance between manifold centroids sharply increased (Fig. 2C, left; paired t-test: stim-on [M = 12.19, SD = 2.40] versus stim-off [M = 3.25, SD = 0.44], t(9) = 11.52, *p* < 0.001, 95% CI [7.2, 10.7], Cohen’s d = 3.64, N = 10), reaching its maximum around 2 s. Simultaneously, normalized intra-cluster variance, quantified by the variance-to-distance ratio, significantly decreased (Fig. 2C, right; paired t-test: stim-on [M = 0.14, SD = 0.03] versus stim-off [M = 0.26, SD = 0.04], t(9) = −12.91, *p* < 0.001, 95% CI [−0.14, −0.10], Cohen’s d = −4.08, N = 10). This concurrent increase in inter-cluster separation (a ‘push’ effect) and decrease in intra-cluster dispersion (a ‘pull’ effect) significantly improved the separability of stimulus categories.

To quantify the efficiency of stimulus encoding in mouse V1, we employed a linear SVM decoder to assess whether the neural manifold geometry in the neural space allowed linear separation of all possible stimulus categorizations. These included three linear separable categorizations in the stimulus feature space (deep versus shallow, narrow versus wide, and spiky versus stubby) and one linearly inseparable categorization (the XOR categorization involving stimulus categories located at two diagonal corners). We found that all four categorizations were linearly separable in the neural space of mouse V1, achieving an overall mean decoding accuracy of 84.5±2.53% across 10 mice (Fig. 2D and Sup. Fig. 1). Specifically, three orthogonal SVM decoding axes emerged in the manifold geometry (Sup. Fig. 2), two corresponding directly to the original stimulus features, and a third representing the XOR distinction not inherently present in stimulus feature space. This indicates that V1 population responses did not merely reflect stimulus features passively but actively generated a new representational dimension to efficiently encode the stimuli (Ma et al., 2025).

Despite the generation of this additional representational dimension, efficient coding remained constrained in a low-dimensional neural space. To illustrate this, we estimated the effective dimensionality (ED) of manifold, a measure of how broadly neural responses are distributed across different directions in neural space, where lower ED indicates that fewer dimensions are sufficient for linear separability (Chung et al., 2018). Immediately following stimulus onset (stim-on), neural manifolds underwent rapid dimensional compression (Fig. 2E; paired t-test: stim-on [M = 4.6, SD = 0.35] versus stim-off [M = 7.1, SD = 0.41], t(9) = −13.9, *p* < 0.001, 95% CI [−2.93, −2.11], Cohen’s d = −4.38, N = 10). Thus, stimulus presence effectively concentrated shared neural variability into a smaller number of modes aligned with stimulus features, yielding a structured, more easily decodable population code (Churchland et al., 2010; Cohen et al., 2020; Jiang et al., 2024; Panzeri et al., 2022; Stringer et al., 2019).

### Stimulus-driven critical state transition at mesoscale level

Having confirmed efficient coding at the macroscale in mouse V1, we next employed pairwise MaxEnt models to investigate its underlying neural basis by examining how sensory inputs reshape pairwise interactions at the microscale. To this end, we separately fitted pairwise MaxEnt models to neuronal population activity recorded during stim-on and stim-off epochs. These models characterize the probability distribution of collective network states constrained by empirically observed first- and second-order statistics (Ganmor et al., 2011). Specifically, we first coarse-grained continuous calcium traces into binary state variables *s_i_* ∈ {−1, 1}, representing whether each neuron exhibited high activity (above its mean, *s_i_* = 1) or low activity (below its mean, *s_i_* = −1) at each time point (Fig. 3A). Such binarization robustly captures emergent dynamics, enabling statistical mechanics approaches to effectively describe continuous neural recordings (Ezaki et al., 2017; Liu et al., 2025; Rosch et al., 2024). We optimized two model parameters, local biases *h_i_* for each neuron and pairwise coupling weights *J*_*ij*_ between neurons (Fig. 3B), via gradient descent to accurately reproduce empirically mean activity (Sup. Fig. 3) and pairwise correlations (Sup. Fig. 4; Methods). The MaxEnt model defines an energy function for binary activity patterns as *H* = − ∑_*i,j*_ *J*_*ij*_*s*_*i*_*s*_*j*_ − ∑_*i*_ *h_i_s*_*i*_, yielding a Boltzmann distribution consistent with empirical first- and second-order neural statistics: *P*(*s*) = *e*^−*H*(*s*)/*T*^/*Z*. Here, temperature T serves as an inverse scaling factor for interaction strengths, with the network’s operational (working) temperature set to *T* = 1, analogous to varying levels of noise.

**Figure 3.**
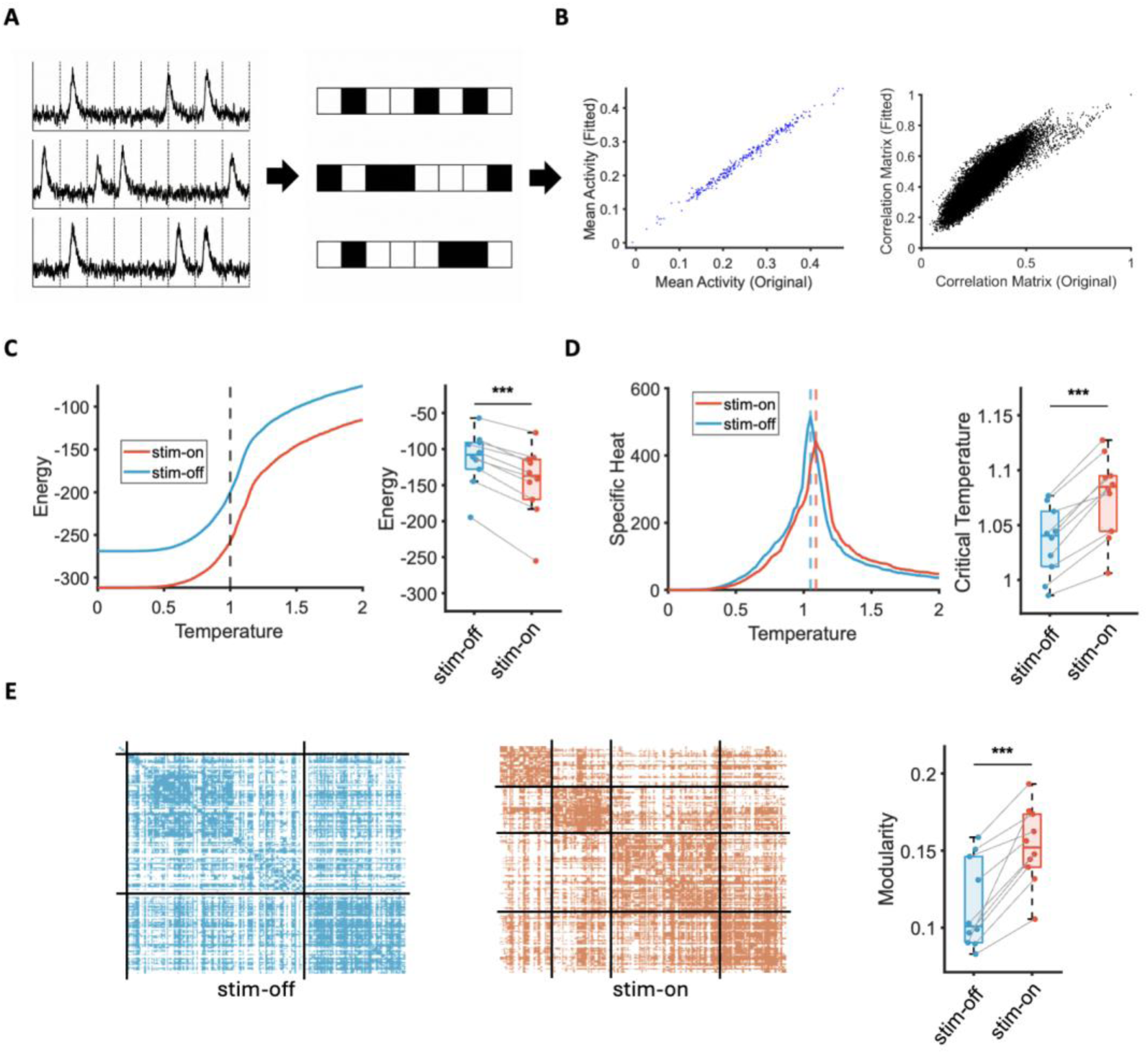
Stimulus-driven changes in collective neuronal interactions and network modularity. (A) Schematic of the neural population state analysis. Calcium traces from neuronal activity during distinct epochs (stim-on and stim-off) were binarized and used to fit pairwise MaxEnt models with matched neuron counts, yielding interaction matrices and neuron-specific biases. The fitted models were then simulated across a range of temperatures to probe network state changes. (B) Comparison between empirical data and fitted network parameters from an example MaxEnt model. Left: Mean neuronal activity (each data point represents one neuron). Right: Pairwise neuronal correlations (each data point represents one neuron pair). (C) Left: Energy curves of MaxEnt models fitted to stim-on (red) and stim-off (blue) epochs across varying temperatures. The dashed vertical line marks the working temperature (*T* = 1). Right: Energy levels at working temperature for stim-off and stim-on epochs across 10 mice. (D) Left: Specific heat curves derived from pairwise MaxEnt models fitted separately to stim-on (red) and stim-off (blue) epochs, plotted across a temperature range. Dashed vertical lines indicate peaks corresponding to critical temperatures. Right: Comparison of critical temperatures from stim-off and stim-on MaxEnt models across 10 mice. (E) Left: Adjacency matrices derived from correlation matrices separated into modular communities using the Louvain algorithm. Modules are visualized during stim-on and stim-off epochs from an example mouse. Within each matrix, neurons are reordered independently based on their community assignments, grouping neurons from the same community together. Right: Modularity measures of functional connectivity networks during stim-off and stim-on epochs across 10 mice. ***: *p* < 0.001.

**Figure 4.**
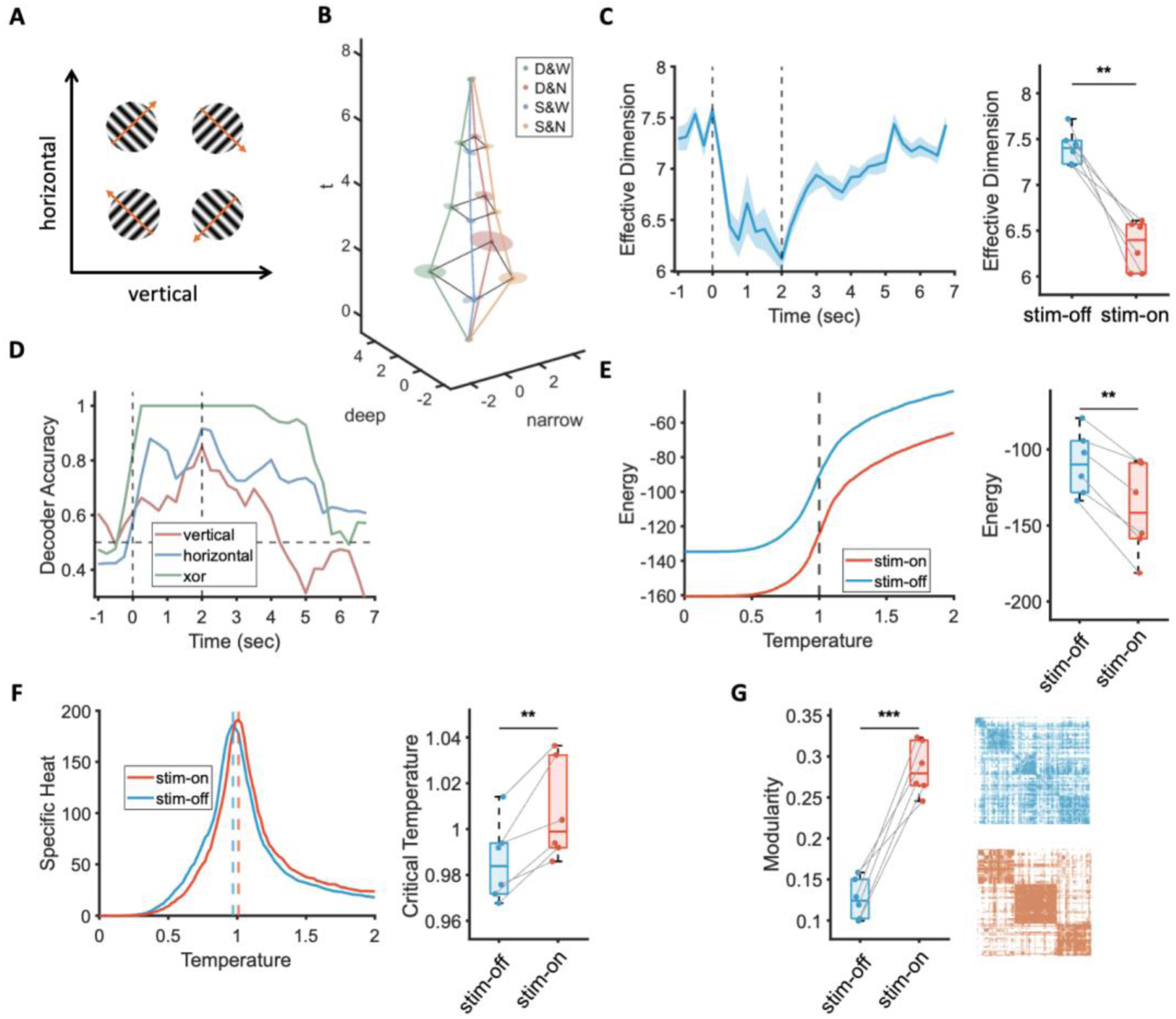
Geometric reorganization and thermodynamic shifts driven by motion stimuli. (A) Schematic of the 2×2 stimulus space, defined by vertical and horizontal motion dimensions, resulting in four distinct motion stimuli. Each stimulus was presented ten times in a randomized sequence. (B) Temporal evolution of neural population responses to the four motion categories averaged over trials in an example mouse. Abbreviations: L, left; R, right; U, up; D, down. Point clouds represent population responses at time points 0, 2, 4, 6, and 8 s relative to stimulus onset, projected onto the two motion-feature axes. Shaded regions indicate ±SD across the 10 trials within each category. (C) Left: Temporal dynamics of effective dimensionality of neural manifolds of an example mouse. Shaded region indicates ±SD from 10 neuron-sampling shuffles. Dashed vertical lines mark stimulus duration. Right: Effective dimensionality measured immediately before stimulus onset (stim-off) and immediately before stimulus offset (stim-on). Values represent averages across all stimuli and trials for six mice. (D) Temporal dynamics of SVM classification accuracy for three pairwise distinctions in an example mouse. Each feature axis indicates pairwise classifications along that dimension. The dashed horizontal line denotes baseline (chance) accuracy (50%). (E) Left: Energy curves of MaxEnt models fitted to stim-on (red) and stim-off (blue) epochs across varying temperatures. The dashed vertical line marks the working temperature (*T* = 1). Right: Energy levels at working temperature for stim-off and stim-on epochs across six mice. (F) Left: Specific heat curves derived from stim-on (red) and stim-off (blue) MaxEnt models across a temperature range. Dashed vertical lines indicate peaks corresponding to critical temperatures. Right: Comparison of critical temperatures from stim-off and stim-on MaxEnt models across six mice. (G) Left: Modularity measures of functional connectivity network during stim-off and stim-on models across six mice; Right: Adjacency matrices derived from correlation matrices separated into modular communities using the Louvain algorithm, shown for stim-off (top) and stim-on (bottom) epochs from an example mouse. Within each matrix, neurons are reordered independently based on their community assignments. ***: *p* < 0.001, **: *p* < 0.01.

To characterize collective-state transitions in the fitted MaxEnt models, we simulated each model across a range of temperatures (*T* ∈ [0,2]) and computed the mean energy *E*(*T*) and specific heat *C*(*T*) = *Var*(*E*)/*T*^2^. As temperature *T* increased, the mean energy *E*(*T*) increased monotonically, with the steepest increase occurring within a narrow transition-like range (Fig. 3C). Within this same interval, energy fluctuations peaked, yielding a pronounced maximum in specific heat *C*(*T*) at critical temperature *T_c_* (Fig. 3D). Importantly, the stim-off model showed a specific heat peaked near the network’s working temperature (*T* = 1), consistent with near-critical dynamics observed during spontaneous activity (Chen et al., 2021; Haimovici et al., 2013; Liu et al., 2025; Nonnenmacher et al., 2017; Tkačik et al., 2015). Here, we interpreted the V1 network state as an equilibrium state, with specific heat divergence serving as a hallmark of criticality. In contrast, the stim-on model showed significantly lower energy at the working temperature (Fig. 3C; paired t-test: stim-on [M = −145.2, SD = 49.10] versus stim-off [M = −112.7, SD = 37.44]; t(9) = −8.22, p < 0.001, 95% CI [−41.5, −23.6], Cohen’s d = −2.60, N = 10), indicating a transition toward a more energetically stable network state.

For the specific heat, the curve obtained from the stim-on model shifted rightward (Fig. 3D), corresponding to a significant increase in the critical temperature *T_c_* (Sup. Fig. 5; paired t-test: stim-on [M = 1.1, SD = 0.04] versus stim-off [M = 1.0, SD = 0.03]; t(9) = 9.20, *p* < 0.001, 95% CI [0.032, 0.052], Cohen’s d = 2.91, N = 10).

**Figure 5:**
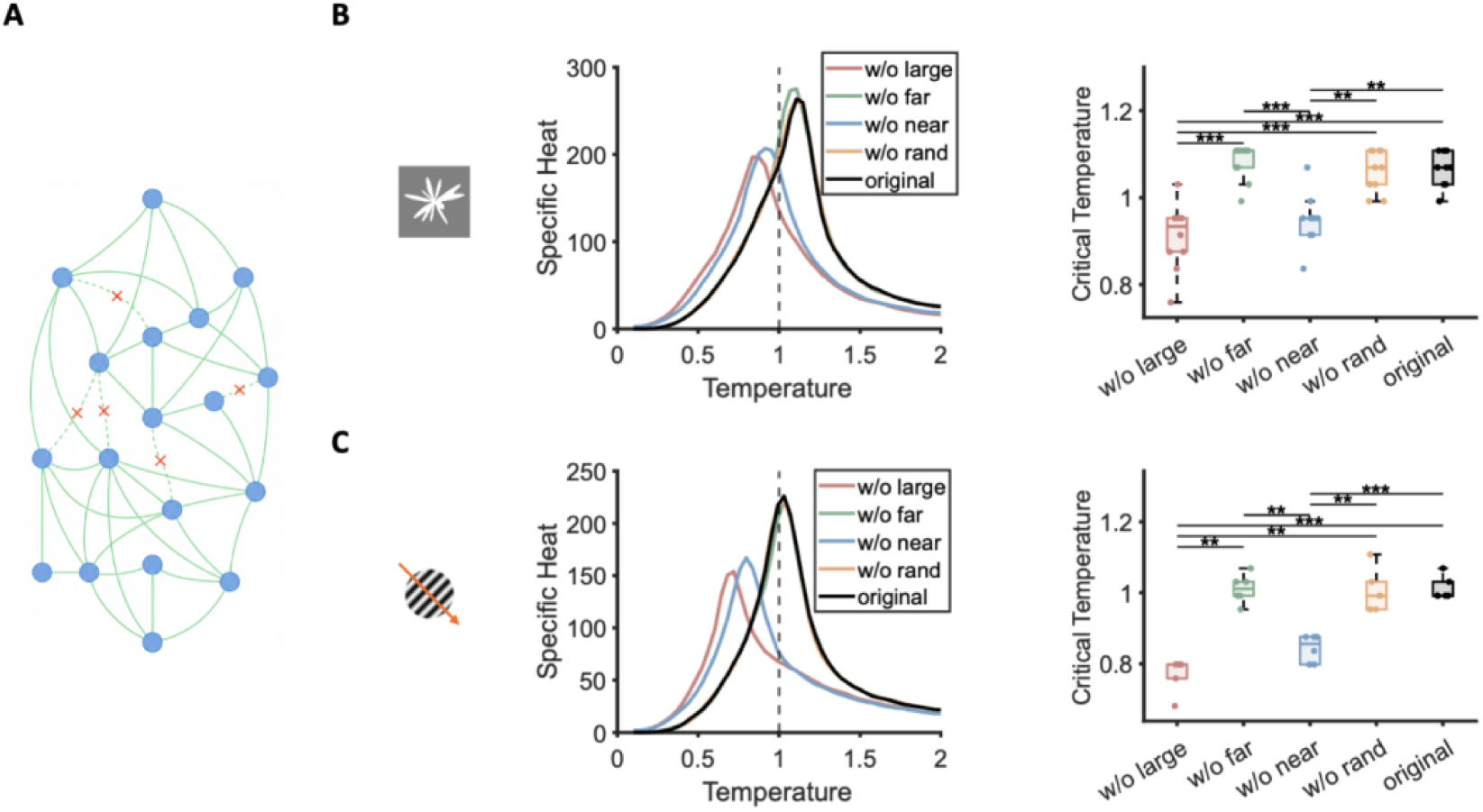
Perturbation Experiments on MaxEnt models. (A) Schematic of perturbations performed on MaxEnt model correlation matrices (J). Selected entries in the fitted correlation matrix were set to zero according to various criteria, effectively removing specific connections and altering direct interactions between neuron pairs. (B) Results of removing an identical number of connections from ten MaxEnt models fitted to the shape experiment, according to four distinct criteria: connections with the largest absolute values (red), connections between neuron pairs with the largest physical distances (green), connections between neuron pairs with the smallest physical distances (blue), and randomly selected connections (yellow). The intact model (black) served as a control. Left: Specific heat curves of these five perturbation models from an example mouse across a range of temperatures. The dashed vertical line indicates the working temperature *T* = 1. Right: Critical temperatures of MaxEnt models under each perturbation averaged across ten mice. (C) Same analyses as in (B), but performed for six mice from the motion experiment.

Given that the network operated at a fixed working temperature (*T* = 1), this increase in critical temperature *T_c_* effectively placed the network state deeper into the ordered, low-temperature regime (i.e., *T* < *T_c_*). In such a regime, entropic fluctuations became suppressed relative to unstructured interactions. Consequently, the stim-on epoch exhibited reduced specific heat at *T* = 1, indicative of fewer accessible thermodynamic states. From this thermodynamic viewpoint, sensory drive biases the network toward structurally ordered states, limiting global fluctuations while preserving structured shared variability within relevant neural subspaces.

Specifically, stimulus presentation broadened the inferred interaction distribution and increased the leading eigenvalues of the interaction matrix (Sup. Figs. 6 and 7), suggesting that the interaction structure became more heterogeneous and dominated by a small set of strong interaction modes. This spectral concentration provided a plausible mechanism for collapsing shared variability into fewer covariance modes, aligning with the observed reduction in effective dimensionality identified in the manifold analysis. To further illustrate this point, we characterized the network topology explicitly by calculating pairwise Pearson correlation distributions across all neuron pairs (Sup. Fig. 8). The Kullback–Leibler (KL) divergence estimated from Kernel Density Estimation (KDE) between stim-on and stim-off correlation distributions was small (0.061±0.047), indicating overall similarity, although mean connectivity strength was slightly higher during stim-on epoch. However, the stim-on epoch exhibited a significantly broader distribution of correlation values (paired t-test: stim-on [M = 0.21, SD = 0.018] versus stim-off [M = 0.17, SD = 0.020]; t(8) = 9.76, *p* < 0.001, 95% CI [0.025, 0.041], Cohen’s d = 3.25, N = 9, excluding one outlier exceeding 3 SD), reflecting increased polarization. As the separation between strong and weak connections widened, contrasts between network clusters became more pronounced (Fig. 3E, left). Correspondingly, adjacency matrices derived from correlation matrices during stim-on epoch exhibited significantly higher modularity (Fig. 3E, right; paired t-test: stim-on [M = 0.15, SD = 0.025] versus stim-off [M = 0.12, SD = 0.029]; t(9) = 6.11, *p* < 0.001, 95% CI [0.024, 0.052], Cohen’s d = 1.93, N = 10).

**Figure 6.**
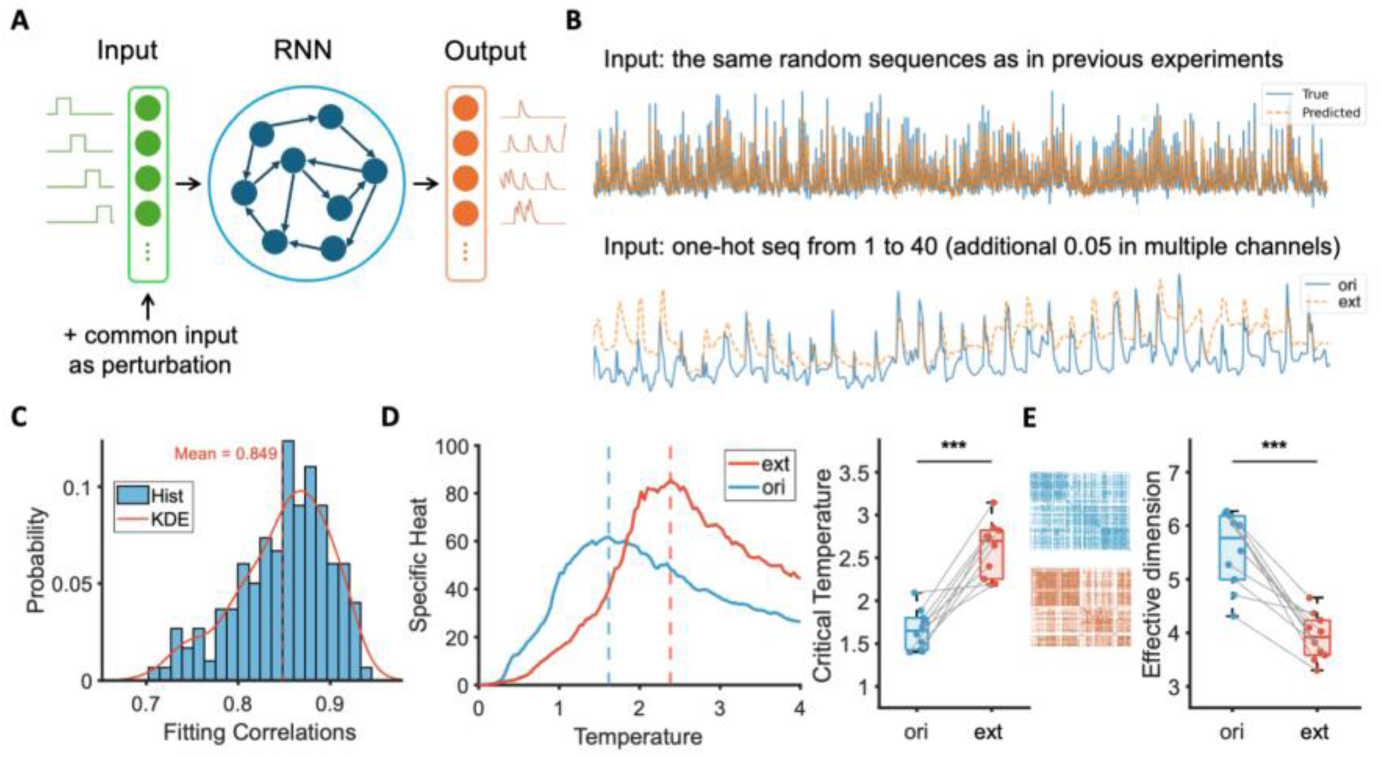
Perturbation experiments on RNN surrogate models. (A) RNN surrogate models were trained to reproduce calcium traces (orange) recorded neurons during the shape experiment, driven by 41-dimensional exemplar-specific input signals (green square waves). (B) Top: Example neuron response when input sequence matches experimental stimulus sequence; observed empirical calcium trace (blue) and RNN-generated prediction (orange). Bottom: Example neuron response illustrating RNN predictions under two distinct input conditions: original sequential inputs (ori; blue) versus the same sequence with additional constant nonzero inputs added to channels without stimulus presentation (ext, orange). (C) Probability distribution of correlation coefficients between all RNN-predicted neuronal responses and empirical data for an example mouse; the red curve indicates KDE. Data correspond to the top panel in (B). (D) Left: Specific-heat curves derived from MaxEnt models fitted to RNN responses under the original (ori) and additional external input (ext) conditions, across a range of temperatures. Dashed vertical lines indicate critical temperatures. Right: Critical temperatures of MaxEnt models fitted to neuronal responses from the ten RNN models under both input conditions. (E) Left: Visualization of modular structures of network responses under ori and ext input conditions; Right: Effective dimensionality of neural manifolds immediately before stimulus offset under two input conditions averaged across the 10 RNN surrogate models. ***: *p* < 0.001.

Thus, neuronal interactions became increasingly modular during stimulus presentation, characterized by strengthened within-module correlations and reduced between-module correlations. Consequently, sensory inputs did not produce uniform decorrelation; instead, they polarized the correlation structure, effectively localizing shared variability within specific modules and restricting global fluctuations to a smaller number of dominant modes. This modular reorganization provides a plausible mesoscale mechanism underlying macroscale efficient coding, facilitating stimulus-aligned manifold compression and linear decoding performance.

### Replication of Thermodynamic Shifts and Geometric Reorganization with motion stimuli

Having established thermodynamic shifts and geometric reorganization with shape stimuli, we further tested the generality of these findings using motion stimuli in a subset of six mice from the same previously described cohort (N=10). Black-and-white moving gratings, generated using sine-wave functions, were defined by two independent spatial dimensions: horizontal motion (left vs. right) and vertical motion (up vs. down). Combining these two orthogonal spatial dimensions resulted in four distinct stimulus conditions: upper-left, lower-left, upper-right, and lower-right (Fig. 4A). The spatial and temporal frequencies of these gratings were selected to match the known preference range of mouse V1 neurons (Andermann et al., 2011). The experimental paradigm was identical to that used in the shape experiment. For data analysis, since gratings inherently lacked within-category exemplary variability, individual trials were treated as distinct samples. Consequently, the ten repeated trials per stimulus provided distinct responses, enabling the generation of point clouds directly comparable to those in the shape experiment.

Motion stimuli elicited geometric reorganization similar to that observed with shape stimuli (Fig. 4B), revealing a clear signature of efficient coding at the macroscale. Projecting trial-averaged neural activity patterns corresponding to the four motion directions into state space resulted in four distinct and well-separated clusters (Fig. 4B). Neural manifolds showed similar temporal dynamics, with inter-cluster distance significantly increased (paired t-test: stim-on [M = 6.8, SD = 1.03] versus stim-off [M = 2.3, SD = 0.43]; t(5) = 8.57, p < 0.001, 95% CI [3.16, 5.87], Cohen’s d = 3.50, N = 6), accompanied by a reduction in normalized intra-cluster variance (paired t-test: stim-on [M = 0.26, SD = 0.023] versus stim-off [M = 0.42, SD = 0.078]; t(5) = −4.45, p = 0.007, 95% CI [−0.262, −0.070], Cohen’s d = −1.81, N = 6). The effective dimensionality rapidly decreased following motion stimulus onset (Fig. 4C; paired t-test: stim-on [M = 6.34, SD = 0.27] versus stim-off [M = 7.41, SD = 0.19]; t(5) = −6.82, p = 0.001, 95% CI [−1.472, −0.666], Cohen’s d = −2.79, N = 6). Additionally, pairwise linear SVM decoders trained to discriminate horizontal (left vs. right), vertical (up vs. down), and XOR stimulus categories demonstrated consistently high decoding performance (Fig. 4D and Sup. Fig. 2), confirming that the stimulus geometry defined by motion features was also reliably preserved. Together, the observed dimensional compression and improved linear decoding performance indicate that motion stimuli similarly constrained V1 dynamics onto a lower-dimensional manifold aligned with stimulus features, thereby facilitating efficient neural coding.

Similar results have been obtained at the mesoscale. MaxEnt models for neural responses to motion stimuli showed lower energy at the network’s working temperature (*T* = 1) during the stim-on epoch compared to stim-off epoch (Fig. 4E, paired t-test: stim-on [M = −140.0, SD = 29.7] versus stim-off [M = −109.3, SD = 20.9]; t(5) = −6.23, *p* = 0.002, 95% CI [−43.3, −18.0], Cohen’s d = −2.54, N = 6).

Additionally, the specific heat peak shifted rightward during the stim-on epoch, corresponding to a significantly increased critical temperature (Fig. 4F, paired t-test: stim-on [M = 1.007, SD = 0.022] versus stim-off [M = 0.986, SD = 0.017]; t(5) = 4.72, *p* = 0.005, 95% CI [0.010, 0.033], Cohen’s d = 1.93, N = 6). Furthermore, the presentation of motion stimuli broadened the inferred interaction distribution (Sup Fig. 6) and increased the leading eigenvalues of the interaction matrix (Sup. Fig. 7), indicating that motion stimuli similarly enhanced local modular structure, shifting the network to a more ordered and energetically stabilized regime. Consistent with this, the correlation distribution in the stim-on epoch was significantly broader (paired t-test: stim-on [M = 0.329, SD = 0.037] versus stim-off [M = 0.172, SD = 0.018]; t(5) = 18.63, *p* < 0.001, 95% CI [0.14, 0.18], Cohen’s d = 7.61, N = 6). Network modularity also increased significantly during the stim-on epoch (Fig. 4G; paired t-test: stim-on [M = 0.285, SD = 0.032] versus. stim-off [M = 0.126, SD = 0.024]; t(5) = 9.42, *p* < 0.001, 95% CI [0.12, 0.21], Cohen’s d = 3.85, N = 6).

Together, these findings from the motion experiment replicated those observed in the shape experiment, suggesting that stimulus-driven geometric reorganization and thermodynamic shifts are not specific to particular stimulus features. Instead, they may represent a general property of neuronal population in mouse V1, where sensory inputs stabilize neural dynamics onto efficient manifolds through reorganization of microscale neuronal connectivity.

### A causal link between thermodynamics and geometry

Across both the shape and motion experiments, sensory inputs consistently produced multiscale effects in mouse V1, including a shift of the specific heat peak toward a more ordered regime, reorganization of functional connectivity into increasingly modular subnetworks, and dimensionality compression accompanied by improved linear decoding performance. These coincident changes appear to reflect a coordinated causal cascade rather than isolated responses to sensory inputs. Indeed, under a weakly magnetized mean-field approximation, collective neural activity is controlled by the dominant spectral mode of the coupling (interaction) matrix, which sets the critical temperature of the network. This interaction spectrum also shapes population variability: in the linear-response regime, each collective mode contributes fluctuations that grow as the system approaches criticality, with the leading mode amplifying most strongly. When variability is modest, population responses occupy an approximately ellipsoidal manifold whose principal axes align with these eigenmodes. The effective dimensionality of this manifold is set by how variance is distributed across modes, decreasing as variability becomes concentrated into fewer dominant directions. Thus, stimulus-dependent reconfiguration of the interactions that strengthens the dominant collective mode can both raise the critical temperature and concentrate fluctuations within a low-dimensional subspace, providing a direct analytical link among microscale neuronal couplings, mesoscale thermodynamic states, and macroscale manifold geometry (for mathematical derivations, see Appendix).

To demonstrate the influence of the eigenvalue spectrum on the critical temperature, we first selectively perturbed the previously fitted correlation matrix *J* (Fig. 5A) to isolate effects of noise and external fields. Specifically, we removed only 0.2% of the connections from the original *J* based on four distinct criteria: (i) connections with the largest absolute values, (ii) connections between neuron pairs with the largest physical distances, (iii) connections between neuron pairs with the smallest physical distances, and (iv) randomly selected connections. This minimal perturbation preserved overall activity statistics while selectively reducing strong and/or short-range connections, given that strongly interacting neuron pairs typically involve short physical distances. In the shape experiment, removing either the strongest connections or nearest-neighbor connections significantly lowered the critical temperature (Fig. 5B; ANOVA: F(4, 36) = 34.83, *p* < 0.001). In contrast, removing the longest-range or randomly chosen connections produced no reliable effect (Bonferroni-corrected *post-hoc* comparisons: w/o strongest [M=0.910, SD=0.076] versus w/o furthest [M=1.081, SD=0.041]: t(9) = −8.12, *p* < 0.001; w/o strongest versus w/o random [M=1.062, SD=0.048): t(9) = −6.65, *p* < 0.001; w/o strongest versus original [M=1.069, SD=0.041): t(9) = −7.50, *p* < 0.001; w/o nearest [M=0.949, SD=0.059] versus w/o furthest: t(9) = −6.28, *p* < 0.001;w/o nearest versus w/o random: t(9) = −4.79, *p* = 0.006; w/o nearest versus original: t(9) = −5.47, *p* = 0.002). Similar results were observed for the motion experiments (Fig. 5C, ANOVA: F(4, 20) = 71.28, *p* < 0.001). Specifically, removing the strongest or nearest-neighbor connections again produced significant reductions in critical temperature (Bonferroni-corrected *post-hoc* comparisons: w/o strongest [M=0.772, SD=0.047] versus w/o furthest [M=1.011, SD=0.041]: t(5) = −8.23, *p* = 0.003; w/o strongest versus w/o random [M=1.005, SD=0.058]: t(5) = −9.48, *p* = 0.001; w/o strongest versus original [M=1.011, SD=0.032]: t(5) = −11.35, *p* < 0.001; w/o nearest [M=0.843, SD=0.038] versus w/o furthest: t(5) = −8.76, *p* = 0.002; w/o nearest versus w/o random: t(5) = −8.72, *p* = 0.002; w/o nearest versus original: t(5) = −10.28, *p* < 0.001). Thus, pruning these large connections reduced the variance of the correlation distribution and attenuated the leading principal components, thereby shifting the network toward a more disordered regime.

Second, we employed a recurrent neural network (RNN) surrogate model to further illustrate how changes in thermodynamic states affect the geometry of neural manifolds, an aspect unable to be directly demonstrated using MaxEnt models alone. Specifically, for each mouse in the shape experiment, we trained a gated recurrent unit (GRU) network to predict calcium activity recorded from all R&R neurons. The stimulus sequence, comprising 40 exemplars spanning four categories plus the gray-screen baseline representing spontaneous activity, was represented as a 41-dimensional, one-hot encoded input vector (Fig. 6A). The RNN surrogate model was trained to reproduce the corresponding neuronal calcium traces (Fig. 6B, top). Rather than performing a simple stimulus-classification task, the model was explicitly trained to predict the full multivariate neuronal response, thus ensuring that it captured underlying population dynamics as accurately as possible. After training, model predictions showed strong agreement with empirical neural responses (r = 0.83 ± 0.01; mean ± SD across neurons per mouse; Fig. 6C and Sup. Fig. 9). Due to the limited stimulus set used in the motion experiment, we restricted these RNN analyses exclusively to the shape experiment.

Two different stimulus sequences were presented as inputs to the trained model (Fig. 6B, bottom). The first input condition, labeled ‘ori’ (original), consisted of the distinct one-hot encoded inputs (stimuli 1–40) identical to those used during training, thus replicating the empirical network states. The second condition, labeled ‘ext’ (additional external input), introduced a continuous background activation of 0.05 to input channels except the one stimulus presentation. This ‘ext’ condition simulated a common external drive, effectively enhancing external fields and suppressing thermodynamic fluctuations. MaxEnt models fitted on RNN-generated outputs allowed estimation of thermodynamic parameters. Compared to ‘ori’ condition, ‘ext’ condition significantly elevated the critical temperature (Fig. 6D; paired t-test: ext [M = 2.60, SD = 0.32] versus ori [M = 1.66, SD = 0.23]; t(9) = 6.96, *p* < 0.001, 95% CI [0.64, 1.25], Cohen’s d = 2.20, N = 10), indicating that the network shifted toward a more ordered regime and confirming the efficacy of the manipulation. We further quantified the effective dimensionality of neural manifolds constructed by RNN-generated outputs. As predicted, the ‘ext’ condition consistently resulted in significantly decreased effective dimensionality compared to the ‘ori’ condition (Fig. 6E, paired t-test: ext [M = 3.92, SD = 0.43] versus ori [M = 5.55, SD = 0.71]; t(9) = −6.67, *p* < 0.001, 95% CI [−2.18, −1.08], Cohen’s d = −2.11, N = 10), thus replicating the thermodynamic–geometric relationship observed experimentally *in vivo*.

Together, perturbation experiments on coupling strengths within MaxEnt models and input manipulations within RNN surrogate models show that stimulus-driven reweighting of effective neuronal couplings at the microscale can shift the network’s critical temperature at the mesoscale and then reorganize neural manifold geometry for efficient coding at the macroscale. Coupled with mathematical derivations linking the eigenvalue spectrum of the correlation matrix to covariance structure and effective dimensionality, these perturbation results support a causal cascade that begins with changes in coupling structure, leads to shifts in thermodynamic regimes, and ultimately results in dimensional compression of neural responses onto stimulus-aligned manifolds, thereby enabling more efficient coding.

## Discussion

This study addresses a fundamental mechanistic question in systems neuroscience: How do microscale neuronal interactions shape macroscale population coding geometry during sensory processing? Our study links microscale changes in pairwise neuronal couplings to macroscale alternations in the dimensionality and geometric structure of population responses through mesoscopic dynamics of criticality.

Specifically, sensory inputs reorganized effective neuronal couplings within mouse V1 along a thermodynamic continuum, transitioning the network from near-criticality toward a more ordered regime. This transition induced dimensional compression of neural responses onto a lower-dimensional manifold aligned with stimulus features, thereby enhancing coding efficiency by reducing unnecessary variability and emphasizing stimulus-relevant dimensions. By integrating frameworks from criticality and efficient coding, our study supports a unified perspective of the brain as both a thermodynamic and computational system capable of dynamically shifting operational regimes and reconfiguring connectivity patterns in real-time to optimally balance flexibility and precision in representing the world.

Previous studies on the critical brain hypothesis have shown that cortical networks, in the absence of external stimuli, operate near a critical regime characterized by heightened sensitivity to weak inputs and extensive intrinsic variability (Arviv et al., 2015; Liu et al., 2025; O’Byrne & Jerbi, 2022; Shew & Plenz, 2013; Toker et al., 2022). Upon stimulus presentation, however, cortical networks shift toward a more stable regime, characterized by reduced trial-to-trial variability and reshaped noise correlations (Ecker et al., 2010; Ioffe & Ii, 2017; Klar et al., 2025; Ohiorhenuan et al., 2010; Ponce-Alvarez et al., 2015). By directly contrasting critical temperatures derived from pairwise MaxEnt models between spontaneous and stimulus-driven states, our study extends this framework by demonstrating that sensory inputs can actively drive such transition in a controlled and reversible manner. Specifically, sensory inputs selectively increased the variance of local neuronal couplings, thereby shifting the network’s critical temperature *T_c_* further above the operational point. Consequently, the network transitioned into a more stable, structured state marked by greater global sparsity, reflected in fewer but more stable dynamic patterns and reduced energy and specific heat. Importantly, this stimulus-driven transition was rapid and context-dependent, modulating the network’s proximity to criticality in real-time. This capability allows cortical networks to dynamically balance exploratory states during spontaneous activity with stable, precise information representation upon stimulus presentation, without relying on slower processes such as synaptic plasticity or structural rewiring.

Context-dependent structural reorganization within the V1 network initiated a multiscale cascade that ultimately enhanced the efficiency of population coding. Previous studies in visual cortex have indicated that spontaneous neuronal fluctuations are not homogeneous; sensory inputs further reduce shared noise correlations, resulting in more structured neuronal responses (Ecker et al., 2011; Koren et al., 2020; Panzeri et al., 2022), and facilitate the formation of specialized subnetworks that support highly informative population codes (Nigam et al., 2019). Consistent with these findings, we found that the microscale adjustments in effective neuronal couplings promoted the formation of distinct modular communities and increased modularity, effectively localizing shared variability within subnetworks.

Furthermore, sensory inputs selectively strengthened particular subnetworks while simultaneously suppressing others, resulting in a polarized correlation matrix characterized by enhanced intra-cluster connections and reduced inter-cluster connections. By compartmentalizing variability into specific modules, the network effectively reduced global fluctuations, thus maintaining stability while retaining adaptive flexibility. Therefore, this modular reorganization promoted efficient processing of distinct stimulus features, ensuring robust signal transmission within modules and minimizing interference among modules, ultimately leading to a stable and efficient coding architecture.

Consequently, at the macroscale of representational geometry, population activity in mouse V1 collapsed onto a low-dimensional manifold aligned closely with stimulus-feature related dimensions. This dimensional compression led to increased linear separability of neural representations, a hallmark of efficient coding that allows neural networks to exact more meaningful information from stimuli (Baddeley et al., 1997; Barlow, 2012; Bernardi et al., 2020; Bolaños et al., 2024; Chung & Abbott, 2021; Cohen et al., 2020; Gallego et al., 2018; Olshausen & Field, 1996; Stringer et al., 2019). From an information-theoretic perspective, the network selectively eliminated redundant dimensions while preserving variance along stimulus-feature relevant axes that carried maximal informational content. In principle, this dimension reduction could improve the mutual information transmitted per neuronal spike, thereby contributing to more efficient communication across the neural population (Dadarlat & Stryker, 2017; Harris & Thiele, 2011; McGinley et al., 2015; Mitchell et al., 2007; Młynarski & Hermundstad, 2021; Thiele & Bellgrove, 2018). Importantly, this compression of representational space was strongly influenced by the fine structure of neuronal correlations, determining whether informational content within a population continues to grow or reaches a saturation point as additional neurons are added (Kafashan et al., 2021; Moreno-Bote et al., 2014).

Previous studies have identified several factors that modulate the correlation structure within neural networks, including internal brain-state changes associated with attention, adaptation, learning, or neuromodulatory drive (Cohen & Maunsell, 2009; Gutnisky & Dragoi, 2008; Minces et al., 2017; Ruff & Cohen, 2014). In addition, long-lasting correlated codes are often observed in higher association cortices, where strong neuronal coupling and stable correlations across time support robust representations and enhance behavioral readout (Panzeri et al., 2022; Runyan et al., 2017; Valente et al., 2021). Our findings supplement this perspective by showing that sensory inputs themselves could rapidly reshape neuronal couplings of mouse V1 on short timescales, producing stimulus-locked, modular correlation structures that compressed representational manifolds independently of internal state changes such as attention and learning. Further mathematical derivations and perturbation experiments established a causal link showing that selectively strengthening specific network interactions, particularly strong, local neuronal couplings, elevated the critical threshold and concentrated neural variability into fewer dimensions, thereby increasing coding efficiency. This finding supports our hypothesis that the architecture of neuronal couplings determines the collective regime of neural networks, which in turn governs coding efficiency. Consequently, our study provides a unified conceptual framework bridging statistical physics, network science, and information theory, offering a coherent explanation of how sensory inputs dynamically drive efficient coding in the brain.

In conclusion, the coupling-criticality-coding triad identified in this study provides a novel framework for understanding how neural networks balance flexibility and fidelity during sensory processing. However, several important questions remain unresolved. First, the precise biological circuits responsible for the observed changes in neuronal couplings need to be identified. Our current recordings focused exclusively on excitatory neurons within cortical layer 2/3, yet inhibitory interneurons are known to significantly modulate network dynamics, including gain regulation and reduction of neural correlations (Harris & Thiele, 2011; Tunstall et al., 2002). Future work combining data from multiple cell types and cortical layers, particularly focusing on inhibitory circuits and translaminar interactions, will be essential for establishing the generalizability of the coupling–criticality–coding triad across diverse neural networks. Second, although our study focused primarily on bottom-up sensory inputs in mouse V1 by excluding explicit task demands, the exact computational and behavioral advantages of observed shifts in criticality remain speculative without directly assessing cognitive and behavioral outcomes. Future work should explore how top-down factors (e.g., cognitive tasks) or internal states (e.g., arousal) modulate stimulus-driven dynamics, providing a more comprehensive perspective on efficient coding. Finally, our current assessment of criticality relies on an equilibrium thermodynamic description inferred from MaxEnt models, which may oversimplify the true complexity of neuronal interactions. Future work that incorporates more sophisticated models, which account for higher-order dependencies and nonlinear coupling, combined with *in vivo* perturbations (e.g., optogenetic manipulation of specific coupling strengths, would further refine our understanding of efficient coding mechanisms across cortical regions and behavioral contexts.

## Data Availability

All data (neuron traces, trained models, and stimulus timing) and analysis scripts are available on the OSF repository (https://doi.org/10.17605/OSF.IO/3AVPC).

## Author Contributions

J.R.L. and J.L. conceived the study. J.R.L. conducted the imaging experiments, developed the computational models and performed data analysis. L.J. built the data preprocessing system. X.L. established the microsurgical and imaging procedures for mice. J.W. provided equipment and technical support. J.R.L. and J.L. wrote the manuscript.

## Acknowledgements

This work was supported by National Natural Science Foundation of China (T2488101), Beijing Municipal Science & Technology Commission, Administrative Commission of Zhongguancun Science Park (Z221100002722012), and Double First-Class Initiative Funds for Discipline Construction. We thank Xiaojuan Wu for her help with the mouse surgery and methodology section.

## Methods

### Animal preparation and habituation

All animal procedures were performed strictly following the guidelines of Tsinghua University and approved by its Institutional Animal Care and Use Committee (IACUC). Mice were maintained in the Laboratory Animal Research Center at Tsinghua University, Beijing, under controlled conditions (temperature: 22-24°C; humidity: 30-70%; lights on 7:00 AM–7:00 PM).

Experimental mice were generated by crossing Rasgrf2-2A-dCre mice (JAX 022864) with Ai148 (TIT2L-GC6f-ICL-tTA2)-D mice (JAX 030328), resulting in Cre-dependent expression of GCaMP6f (Jackson Laboratory). Seven adult male and three female mice (9-17 weeks old; body weight: 20–30 g) were housed in groups in the specific pathogen-free (SPF) animal facilities. They received intraperitoneal injections of trimethoprim (TMP, 0.25 mg/g; Sigma) on four consecutive days. Food and water were provided ad libitum. Following surgery, mice were individually housed and habituated for at least one week before recordings began.

Mice were habituated to the imaging apparatus for 7 days. On days 1–3, animals freely explored an open-field arena (45 cm × 45 cm) containing the head-fixation stage (30 min per session) and received a food reward after each session. On days 4–6, mice were head-fixed on the stage in the imaging environment for 30 min/day with food reward as positive reinforcement. On day 7, a 30-min pilot imaging session was conducted to assess imaging quality and for inclusion screening.

### Craniotomy operation for wide-field imaging

Mice were anaesthetized with tribromoethanol (Avertin; 1.25%, 0.2 ml per 10 g body weight, intraperitoneal) and maintained with isoflurane (0.5–1.0% in oxygen; 0.5 L/min; RWD Life Science) while body temperature was held at ∼37 °C using a heating pad. The skull was stereotaxically levelled (Bregma–Lambda height difference <0.03 mm; mediolateral <0.03 mm), and a 7 mm circular craniotomy centered 1.3 mm lateral and 2.5 mm anterior to Lambda was performed under a surgical microscope with the dura left intact. A sterile 7 mm coverslip was seated and sealed with tissue adhesive and cyanoacrylate; two skull screws were placed ≥2 mm from the window edge, and a custom aluminum head plate was secured with dental cement with its 10 mm aperture plane parallel to the window. Postoperative care included meloxicam (4 mg/kg, subcutaneous) and ceftriaxone (0.2 ml, intraperitoneal, once daily for 3 days; 80 mg/ml in saline); animals were monitored daily and recovered for at least 1 week before habituation.

### Calcium imaging

Neuronal calcium activity was recorded with the RUSH3D computational mesoscope based on scanning light-field microscopy (Zhang et al., 2024). The system incorporates a custom macro-objective (NA 0.5) and a large-format sCMOS camera (CMV50000, CMOSIS). To surpass the microlens-array sampling limit (69 µm pitch), a piezo tip–tilt actuator implemented a 3 × 3 sub-microlens scan during acquisition, enabling oversampled light-field capture and volumetric imaging with an approximately uniform resolution of ∼2.6 × 2.6 × 6 µm³ over a native field of view of ∼8 × 6 × 0.4 mm³ at 4 Hz. Fluorescence was excited with a 488 nm LED (LDI-488, 89 North) at ∼10% maximum output (∼0.12 mW/mm²). Imaging frames and visual stimuli were synchronized using the Medusa data acquisition system (Bio-Signal Technologies).

Imaging was primarily sensitive to layer 2/3 neurons in cerebral cortex, given the penetration depth of wide-field illumination (∼200–300 µm). GCaMP6f was expressed under the Rasgrf2 promoter, labeling excitatory neurons. As a result, our recordings largely reflect the activity of excitatory pyramidal neurons in layer 2/3 of V1, which represent the primary cortico-cortical output channel and play a central role in hierarchical visual processing. Layer 2/3 pyramidal neurons integrate feature-selective inputs from layer 4 and provide major feedforward output from V1 via long-range projections to higher visual areas (LM, AL and PM) and local projections to deeper layers (including L5), making their activity well suited for probing how sensory signals are transformed into population codes accessible to downstream circuits. In addition, superficial excitatory neurons are optically accessible and yield robust signal-to-noise for large-scale calcium imaging, enabling stable measurements of population dynamics during naturalistic stimulation.

### Fluorescence image processing and neuron extraction

To process the large-scale 3D light-field datasets acquired with the RUSH3D system, we used the standard computational pipeline provided by the system developers (Zhang et al., 2024). The workflow comprised four sequential stages. First, raw light-field measurements were converted into high-resolution spatial–angular images by pixel realignment, following established procedures for scanning light-field microscopy with digital adaptive optics (Lu et al., 2022). Second, to suppress the strong out-of-focus and background fluorescence characteristic of wide-field mouse brain imaging, we applied the multiscale background rejection (MBR) algorithm (Zhang et al., 2021) followed by rigid motion correction performed on the phase-space data using the NoRMCorre algorithm (Pnevmatikakis & Giovannucci, 2017) to stabilize tissue movements. Third, the background-suppressed spatial–angular images were reconstructed into 3D, aberration-corrected volumes using SeReNet (Lu et al., 2025) together with wavefront digital adaptive optics (wDAO) (Wu et al., 2021).

Finally, neuronal regions of interest (ROIs) and corresponding fluorescence time series were extracted using CNMF-E as implemented in CaImAn (Giovannucci et al., 2019). The temporal traces were calculated by Δ*F*/*F0* = (*F − F0)* / *F0*, where *F0* is the mean fluorescence in the ROI averaged over the entire time series, and *F* is the averaged intensity of the ROI.

### Intrinsic signal imaging

We performed intrinsic signal optical imaging (ISOI) over the right visual cortex to obtain retinotopic maps (azimuth and elevation) for visual area delineation and downstream registration. Visual stimuli were generated with a custom Python pipeline and presented on a 60-Hz LCD monitor (1920 × 1080) positioned 15 cm from the eye; screen coordinates were converted to visual-angle coordinates using an arctangent-based spherical correction. Retinotopic mapping used a drifting checkerboard bar sweeping in four directions (bottom-to-top, top-to-bottom, left-to-right, right-to-left; bar width 20°; step size 0.15° per frame; checkerboard contrast updated every 10 frames, ∼6 Hz), with a 2-s pre-gap and 3-s post-gap per direction; the four-direction sequence was repeated 40 times. During imaging, mice were lightly anesthetized with isoflurane (0.5% in oxygen, 0.8 L/min) and maintained at ∼37 °C; wide-field reflectance images were acquired under green (∼525nm) illumination to capture intrinsic hemodynamic signals and surface vasculature, with stimulus presentation and acquisition hardware-synchronized via NI-DAQ.

Retinotopic maps were computed using Fourier phase analysis by extracting, for each pixel, the phase at the stimulus frequency; hemodynamic delay was compensated by subtracting phase maps obtained from opposite sweep directions. Azimuth and elevation phase maps were then combined to compute a visual field sign map from the angular difference between their spatial gradients, and visual areas were delineated from sign reversals following established conventions (Garrett et al., 2014). For atlas-based assignment, imaging data were registered to the Allen Mouse Common Coordinate Framework (CCF, top projection) (Wang et al., 2020) using vasculature-guided transforms: an intrinsic vasculature image was aligned to the CCF, and session-specific fluorescence projection images were registered to the intrinsic vasculature using manually selected vascular control points to estimate a projective transform. Concatenated transforms were applied to neuron footprint centers (with correction for any horizontal flip performed before control-point selection), and each transformed coordinate was assigned to an atlas region by sampling the CCF index map at the corresponding location.

### Experimental protocol

We focused our analysis on the primary visual cortex (V1) of mice, a crucial region for early-stage visual processing. In the shape paradigm, mice were presented with randomized sequences comprising 40 distinct shape stimuli plus a gray screen condition. In the motion-grating paradigm, mice were presented with four drifting-grating directions plus the same blank condition. Each trial consisted of a 2-s stimulus presentation followed by a 6-s interstimulus interval (ISI) of a mean-luminance gray screen. In the shape experiment, each stimulus corresponds to eight trials, while in the motion experiment, there are only 4 stimuli, each corresponding to 10 trials. This design allowed us to systematically investigate the dynamic response properties and adaptability of neuronal networks within V1 to diverse visual inputs. Stimuli were displayed on a 7-inch diagonal screen placed 10 cm in front of the mice. The screen was oriented laterally at an angle of 30° relative to the midline of the mice and tilted downward by an additional 30° to optimally align with their visual field. To minimize simple arousal influence, epochs alternate within a short period of time (8 s here) and the spontaneous activity epoch consisted of a gray screen presentation with the same luminance as in stimulus presentations.

### Stimuli Design

For shape experiment, the visual stimuli consisted of four discrete shape categories defined by two binary factors—’deep vs. shallow’ and ‘narrow vs. wide’. Although conceptually arranged along two orthogonal feature axes for visualization, the design sampled only four categorical points rather than a continuous space. These stimuli were petal-like blobs created by manipulating two main parameters: the number of control points (either 10 for wide or 35 for narrow) and the maximum allowable indentation or minimum control point radius (0% for deep or 50% for shallow) (Frank et al., 2023). Generated patterns correspond to an average 30 degrees in field of view, while the average spatial frequency is close to 0.05 cycles per degree (cpd), chosen to fall near the preferred spatial-frequency range of mouse V1 neurons (Andermann et al., 2011).

Specifically, randomly positioned control points were generated and transformed from Cartesian coordinates into polar coordinates, computing the angular position and radius relative to the shape’s center. To form a closed, continuous boundary, angular coordinates were normalized to a range of 0 to 2π, sorted in ascending order, and the radii reordered correspondingly. This sorting ensured sequential connectivity between control points, preventing boundary overlaps. Subsequently, a Piecewise Cubic Hermite Interpolating Polynomial (PCHIP) was applied to interpolate between sorted polar coordinates, generating intermediate points to produce smooth and continuous shape boundaries. PCHIP interpolation was selected specifically due to its capability to produce smooth, oscillation-free curves that pass precisely through each control point, preserving essential geometric characteristics such as smoothness and gradual curvature transitions.

Stimuli generated via this method were centered within a fixed-size window and presented as white shapes against a uniform gray background. Adjusting the number of control points and minimum control point radius enabled systematic variation along the two axes, resulting in four distinct stimulus conditions: ‘Deep & Wide’ (D&W), ‘Deep & Narrow’ (D&N), ‘Shallow & Wide’ (S&W), and ‘Shallow & Narrow’ (S&N). Specifically, the D&N condition corresponded to spiky shapes, whereas the S&W condition corresponded to stubby shapes based on their angular features. Each condition consisted of 10 unique exemplars, generated by randomizing control point positions and radii, introducing controlled variability within each condition and enabling the characterization of both general and stimulus-specific neural responses in the visual cortex. That is, while all ten exemplars within a given category shared the identical defining feature parameters, they varied due to stochastic variability during generation (e.g., specific locations of vertices and edges), thereby ensuring their independence as distinct visual inputs.

For motion stimuli, we organized conditions in a 2D stimulus plane with two binary factors: upward vs. downward and leftward vs. rightward motions. Movement directions comprised four orientations: upper-right (45°), upper-left (135°), lower-right (315°), and lower-left (225°), forming a balanced 2×2 factorial design. These four directional stimuli, together with a blank stimulus, constituted five conditions, each presented 10 times in a randomized sequence. Gratings were generated using sine function, with gratings presented as white (opacity = 1) on a gray background. Spatial and temporal parameters of the gratings were selected for the highest selectivity on mouse V1 (Andermann et al., 2011); spatial frequency was set to 0.05 cycles per degree, and temporal frequency was set to 1 Hz.

We performed two stimulus paradigms: a shape paradigm and a motion-grating paradigm. The shape paradigm was recorded in 10 mice (10 imaging sessions/experiments). The motion-grating paradigm was recorded in 6 imaging sessions/experiments (6 mice out of 10 mice above).

### Data Preprocessing

All analyses were performed using custom MATLAB scripts (MATLAB R2024b). For each neuron, we first z-scored the ΔF/F trace using the mean and standard deviation computed over the full recording. To identify stimulus-responsive neurons, we defined three time windows relative to stimulus onset: baseline (−1 to 0 s), evoked response (1 to 3 s; accounting for calcium latency), and post-stimulus recovery (3 to 5 s). For each stimulus condition, a neuron was considered responsive if its mean activity in the evoked window exceeded both baseline and recovery in ≥80% of trials and this increase was statistically significant across trials (Wilcoxon signed-rank test, two-sided, α = 0.05). To identify trial-consistent neurons, we defined each neuron’s preferred condition as the stimulus (or stimulus category) that produced the largest mean evoked response. A neuron was considered reliable if its evoked responses to the preferred condition were significantly larger than responses during blank (gray-screen) trials (*p* < 0.05, two-sided t-test across trials) and exceeded an effect-size criterion (Cohen’s d > 1). Neurons satisfying both criteria were termed reliably responsive (R&R) and were used for all subsequent analyses (Marshel et al., 2011).

Neuronal responses of each neuron underwent standardization via z-scoring with its own mean and standard deviation of the fluorescence trace across the whole experiment. To compare stimulus-evoked and spontaneous activity, we segmented each 8-s trial into a stimulus-driven epoch (stim-on) and a post-stimulus epoch (stim-off). Shape paradigm: stim-on = 0–3.5 s; stim-off = 3.5–8 s. Motion paradigm: stim-on = 0–4.5 s; stim-off = 4.5–8 s. The dividing time was chosen based on the decay of the population-averaged calcium response after stimulus offset, such that the stim-on window captures the stimulus-locked response including calcium delay/decay while the stim-off window reflects gray-screen activity closer to baseline. These epoch boundaries were fixed a priori and applied to all mice.

### Manifold Geometry and Effective Dimension

We quantified population-level separation between two diagonal conditions in the 2D feature space (spiky vs. stubby in the shape experiment; upper-left vs. lower-right in the motion experiment) using a subsampling and shuffling procedure applied to trial-averaged neuronal activity. For each time point, we quantified (i) between-condition separation and (ii) within-condition dispersion for the two diagonal conditions (shape: spiky vs. stubby; motion: upper-left vs. lower-right). For shape, the 10 exemplars within each condition were treated as samples after averaging responses across repeated presentations; for motion, the 10 trials were treated as samples. In each of 10 subsampling iterations, we randomly selected 60% of R&R neurons and computed the condition centroids (mean response vectors across samples). Separation was defined as the Euclidean distance between the two centroids. Within-condition dispersion was quantified as follows: for each neuron, we computed the standard deviation of its responses across the 10 samples within each condition, and then averaged this standard deviation across neurons and across the two conditions. Cluster separability was summarized by a variability–distance ratio as normalized variance, defined as this average within-condition variability divided by the Euclidean distance between the corresponding mean population response vectors.

Chung and colleagues (2018) developed a theory of linear separability for object manifolds, which defines the effective manifold dimension and classification capacity using anchor points at the edges of each manifold. Following this framework, we applied their manifold analysis code to trial-averaged population responses for the two diagonal conditions in the 2D feature space, treating repeated samples (10 per condition) as points on each manifold. For each time point, we first randomly subsampled two-thirds of the R&R neurons without replacement and split the resulting response matrix into four condition-specific blocks (corresponding to the four stimulus categories); only the two diagonal conditions were used as manifolds in the analysis. For these two manifolds, we computed the manifold capacity, radius, and effective dimension using the original implementation with zero margin (κ = 0) and 300 randomly sampled directions, and then averaged the effective dimension across the two manifolds. This subsampling and analysis procedure was repeated 10 times, and the reported manifold dimension for a given time point was taken as the mean effective dimension across subsamples. Codes used were directly from the original paper (Chung et al., 2018).

### SVM Classification Analysis

Linear Support Vector Machines (SVMs) were utilized to decode categorical information within neural data and identify distinct neural space dimensions. We segmented trial-averaged neural responses into training and testing sets across various temporal bins spanning from −1s to 7s relative to stimulus onset. Within each time bin, half of the trials were randomly assigned to the training set, with the remaining assigned to the testing set. MATLAB’s fitclinear function was employed, maintaining default hyperparameters but adjusting the regularization parameter lambda to 1, balancing complexity and generalization capacity. Classification accuracy was evaluated over time to generate temporal profiles reflecting the discriminative capability of neural populations. The final decoding accuracy was obtained by first averaging the accuracies from the different binary classifiers and then averaging these values across classifiers.

Note that in the time course of decoder accuracy, another different SVM was trained and tested independently at each time point, using 50% of the data for training and 50% for testing. Because the number of samples at each time point was small (eight trials in total), the resulting estimates are expected to be somewhat unstable.

Moreover, although time 0 is defined as stimulus onset, the recording frequency was relatively low (4 Hz), so the actual onset could occur within ±0.25 s around the nominal 0 time point. Together with the limited number of samples, it is therefore not surprising that SVM decoding accuracy at time 0 occasionally exceeds chance level.

To quantify the similarity between decision boundaries generated by SVM classifiers, we performed angle analysis. Specifically, we computed the angles between n-dimensional unit vectors (β_*i*_ and β_*j*_), using the formula: θ = *arccos*(|β_i_^*T*^β_*j*_|). These vectors represented the decision boundaries derived from SVMs trained on different conditions. Orthogonal decision boundaries (angles approaching 90 degrees) indicate independent feature encoding, implying neural representations optimized for discriminative efficiency. To verify robustness, angle analyses were repeated eight times per classifier, confirming consistency in observed patterns across iterations.

### Pairwise Maximum Entropy Model

We analyzed activity from all R&R neurons using a pairwise maximum entropy model (MEM) (Jaynes, 1957; Schneidman et al., 2006). To define the discrete network states required for this statistical-mechanical framework, we binarized the calcium signal of each neuron *i* at each time point *t* after a bandpass filtering (0.1–1 Hz) (Cramer et al., 2019; Schwalm et al., 2017; Vanni et al., 2017). The lower cutoff frequency (0.1 Hz) was used to remove baseline drifts arising from photobleaching and other slow fluctuations (e.g., gradual changes in physiological state, temperature, or slow drift of the optical path). The upper cutoff frequency (1 Hz) was chosen to attenuate high-frequency measurement noise while, under the constraint of a 4-Hz sampling rate, preserving a reliable representation of the stimulus fundamental frequency (≈0.5 Hz) and its potential low-order harmonics. The signal was assigned a value of +1 (active) if it exceeded its overall mean (0), and −1 (inactive) if it fell below the mean. This constitutes a coarse-graining step, transforming continuous, high-dimensional signals into a more tractable state space capable of revealing collective properties—a theoretically grounded approach (Hopfield, 1984). Such binarization of continuous neuroimaging data (including fMRI and calcium signals), followed by application of the MaxEnt model, has been successfully employed to uncover emergent energy landscapes and functional interactions that are not directly observable from the raw continuous signals themselves (Ezaki et al., 2017; Kandeepan et al., 2020; Rosch et al., 2024). We opted for direct binarization, as spike deconvolution carries multiple limitations for our single-photon widefield imaging data. Previous studies have demonstrated that deconvolution algorithms yield unreliable results under high-background conditions like single-photon despite the subtraction (Rupprecht et al., 2021). Also, different deconvolution algorithms can produce qualitatively divergent outcomes even when applied to the same calcium imaging dataset (Evans et al., 2019), and there is no standardized set of deconvolution parameters for single-photon calcium imaging.

This model describes the probability distribution P(s) of collective neuronal activity patterns, constrained by empirically observed mean activities 〈*s*_*i*_〉 and pairwise correlations 〈*s*_*i*_*s*_*j*_〉, while maximizing entropy in other respects. The neuronal state distribution follows a Boltzmann distribution (Jaynes, 1957):

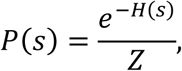

where Z is the normalizing partition function:

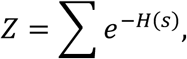

and H is the energy function:

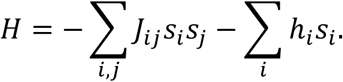

Here, *J*_*ij*_ represents the functional interaction between neuron *i* and neuron *j*, and *h_i_* denotes the activation bias. Here, the parameter J differs from simple, model-free pairwise correlations (functional connectivity); it represents the effective pairwise coupling (also termed as functional interaction) between neurons, which statistically controls for indirect effects mediated by other units in the network (Ganmor et al., 2011). To estimate optimal parameters, we employed a pseudo-likelihood maximization algorithm using gradient descent as in previous research fitting large spin systems (Ezaki et al., 2017; Rosch et al., 2024; Tkačik et al., 2014), addressing the computational challenges posed by large neuron populations (hundreds of neurons):

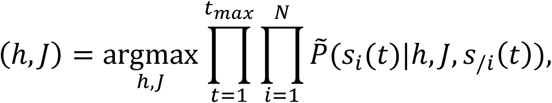

where *P*∽ is the single-neuron Boltzmann conditional distribution given the states of other neurons. To estimate the model’s parameters h and J, we used a gradient descent scheme, which updates the parameters by comparing the empirical mean and correlation values to the predicted mean and correlation values by the model. The update equations are as follows:

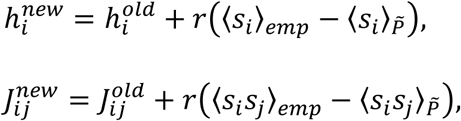

with a learning rate *r* = 0.05. We initialized parameters at zero, enforced *J*_*ij*_ = *J*_*ji*_ and *J*_*ii*_ = 0, and iterated until convergence (change in error < 1e-8). Fit quality was assessed by comparing empirical and model-predicted first- and second-order statistics (Supplementary Figs. 3–4). Model fitting quality for first- and second-order statistics is shown in Sup. Figs. 3 and 4. The statistical simulation results of the model are all strongly positively correlated with the empirical data, and the data points are close to the diagonal line of y = x, indicating the validity of the fit. The superscripts new and old represent the parameters after and before a single updating step. 〈*s*_*i*_〉_*P*∽_ and 〈*s*_*i*_*s*_*j*_〉_*P*∽_ are the mean and correlation with respect to distribution *P*∽:

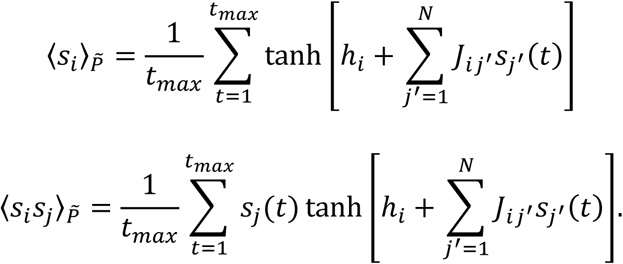

Although the pseudo-likelihood approach employs a mean-field approximation, its estimates converge to maximum-likelihood solutions as the number of time points *t*_*max*_ → ∞ (Besag, 1975). The details of pseudo-likelihood maximization algorithms and scripts used refer to (Ezaki et al., 2017).

### Thermal Characteristics of MaxEnt Model

To thoroughly investigate the modeled neural activation patterns, we utilized Markov Chain Monte Carlo (MCMC) methods with the Metropolis–Hastings algorithm (Hastings, 1970), simulating individual neuronal state transitions and collective neuronal interactions. Temperature effects were incorporated into the Boltzmann distribution as:

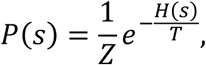

with the partition function Z given by:

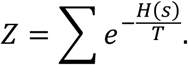

Simulations began with random initial states and included 1,030,000 iterative steps. The initial 30,000 steps were discarded to achieve thermal equilibrium, with subsequent states sampled at intervals of 500 steps to mitigate autocorrelation.

Simulations were performed across decreasing temperatures, from *T* = 2 to *T* = 0, each initialized using the final state of the preceding higher-temperature simulation.

The specific heat and susceptibility were calculated from energy fluctuations using the fluctuation-dissipation theorem (Kubo, 1966):

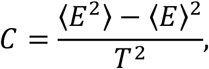

where *E* represents system energy, *T* is the temperature, and angle brackets indicate averages over MEM. For each state, all neurons’ states *s*_*i*_ and network parameters ℎ and *J* are all known, then we can calculate the system energy exactly as the definition. For each fitted MaxEnt model, we repeated the simulation 10 times for robust measurement.

### Modularity Measurements

To assess neuronal modularity, we computed pairwise Pearson correlation matrices of neuronal activity and analyzed community structures using the Louvain modularity algorithm (Blondel et al., 2008). For each experiment and epoch (stim-on and stim-off), we computed an N × N Pearson correlation matrix across R&R neurons using z-scored activity concatenated over all frames belonging to that epoch across all trials.

Negative correlations were set to zero to obtain a non-negative weighted adjacency matrix. Community structure and modularity Q were computed using the Louvain algorithm (resolution parameter γ = 1; weighted version). Because Louvain optimization is stochastic, we repeated community detection 10 times and report the mean modularity across runs. Modularity scores, indicating strength of community structures, were calculated as:

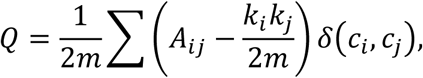

where *A*_*ij*_ represents adjacency matrix elements, *k*_*i*_ and *k*_*j*_ denote node degrees, *m* is the total number of edges, and δ(*c*_*i*_, *c*_*j*_) equals 1 if nodes *i* and *j* belong to the same community, otherwise 0.

### Neural Interaction Perturbation

To investigate the influence of pairwise interactions on neural population dynamics, we additionally fitted a pairwise maximum entropy model to whole calcium traces from identified R&R neurons for each mice, deriving an interaction matrix ***J***. This model-based measurement method has been shown to be more sensitive to structure-driven relationships than the original FC and can serve as an intermediary connecting SC and FC (Ashourvan et al., 2021). Using this fitted matrix ***J***, we simulated neural dynamics employing Markov Chain Monte Carlo (MCMC) methods across varying temperatures to estimate the critical temperature.

We examined the impact of different connectivity-pruning strategies on network dynamics by constructing several variants of the empirical coupling matrix ***J*** defined over the R&R neurons. For each neuron, we extracted its physical coordinates in the imaging plane and computed the full pairwise Euclidean distance matrix between all neuron pairs. Because ***J*** is symmetric, we considered only the upper-triangular (off-diagonal) entries as unique undirected connections and, for each pruning condition, removed a fixed fraction *p* = 0.002 (0.2%) of these pairwise couplings by setting the corresponding *J*_*ij*_ and *J*_*ji*_ elements to zero. Four pruning schemes were implemented: (i) Strongest – removal of the connections with the largest coupling strength; (ii) Furthest – removal of the connections linking the most distant neuron pairs in physical space; (iii) Nearest – removal of the connections between the closest neuron pairs; and (iv) Random – removal of an equally sized, randomly selected subset of connections. In addition, the unmodified connectivity matrix was retained as an original control condition. For each of these five matrices (Original + four pruned variants), we simulated an Ising model with the same fitted external field over temperature values linearly spaced between *T* = 2 and *T* = 0. Each simulation consisted of 2,100,000 iterations, of which the initial 100,000 iterations were discarded as thermalization, and observables were sampled every 500 iterations. From the simulation output we obtained the temperature dependence of the specific heat; for visualization, the resulting curves were smoothed with a three-point moving average, and the temperature corresponding to the peak was used to characterize the effective critical-like behavior of each pruning condition.

### Recurrent neural network

We conducted RNN analysis for shape condition, because in the motion experiment, the dimensionality of the input (four distinct stimuli encoded as four one-hot channels) is far smaller than that of the output (the continuous responses of several hundred neurons). We implemented all recurrent network analyses in Python using PyTorch, based on the same R&R-selective neuron population used in the main analyses. For each recording we used selective Calcium activity traces and the corresponding stimulus identity sequence. The trace matrix had dimensions *N* × *T*, where *N* is the number of selectively responding R&R neurons and *T* is the number of imaging frames. The stimulus array was a length-T vector of integer labels indicating which of 40 visual stimuli or a gray screen was present at each time point. We then transformed the sequence into a one-hot representation with dimensionality *D* = 41 (40 stimuli + gray). This produced an input tensor of shape (*T*, *D*) that preserved the original temporal resolution of the calcium recordings. Each time step consisted of a 41-dimensional stimulus code and a corresponding N-dimensional population response.

The recurrent model was a gated recurrent unit (GRU)–based RNN designed to map the stimulus sequence to the simultaneously recorded R&R responses. The network consisted of two stacked GRU layers followed by a linear readout.

Specifically, the GRU received the D-dimensional one-hot stimulus vector at each time step and produced a hidden state of size *H* = 2*N*. The second GRU layer operated on this hidden state sequence with the same hidden dimensionality, and the terminal hidden representation at each time step was passed through a linear fully connected layer to yield an N-dimensional output vector corresponding to the predicted Calcium activity of each R&R neuron. The forward pass thus produced an output tensor of shape (*T*, *N*) and an intermediate hidden-state tensor (*T*, *H*). By relying on GRUs rather than vanilla RNN units, the model could selectively retain or reset temporal information via update and reset gates, enabling it to capture stimulus-specific temporal integration and decay dynamics.

Model training was formulated as sequence-to-sequence regression from the one-hot stimulus sequence to the continuous calcium traces. We used mean squared error (MSE) between predicted and recorded activity at each time step and neuron as the primary loss. Besides using a modest number of parameters, to prevent overfitting and constrain the connectivity structure, we additionally implemented strong L2 regularization on the recurrent and readout weights (Sussillo et al., 2015). A dedicated routine traversed all trainable parameters and accumulated the squared ℓ2-norm separately for recurrent weights in the GRU layers (regularization coefficient λ_*rnn*_) and the weights in the final fully connected readout layer (coefficient λ_*fc*_). The total training loss was defined as

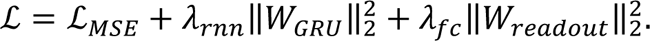

Optimization was performed with the Adam algorithm (learning rate =0.001, β_1_ = 0.9, β_2_ = 0.999, ε = 10^−8^), training on the full continuous sequence over multiple epochs. The network’s hidden state was initialized to zero at the start of each epoch, and gradients were backpropagated through the entire sequence. During training we tracked three quantities across epochs: the main MSE loss, the L2 penalty term, and their sum, allowing us to monitor both fit quality and the contribution of regularization.

After training, we evaluated how well the RNN captured the temporal structure of individual neurons. With the model in evaluation mode and gradients disabled, we passed the entire empirical stimulus sequence through the network once to obtain predicted responses ŷ_*i*_(*t*) for each neuron *i*. For quality assessment, we computed the Pearson correlation coefficient between each neuron’s predicted and recorded time series, *corr*(ŷ_*i*_(*t*), *y*_*i*_(*t*)), and summarized the distribution of correlations across neurons. We also identified the top and bottom subsets of neurons (e.g. best and worst five neurons) according to this correlation metric and visualized their full traces, plotting true and predicted activity on the same time axis to qualitatively assess how well the network captured response amplitude, latency, and temporal modulation.

To probe the network’s dynamics under controlled conditions, we constructed a synthetic test sequence that systematically sampled the stimulus set. This sequence sequentially presented each of the 40 stimuli for 10 consecutive time steps, followed by 40 time steps of the gray screen, and repeated this pattern once, yielding a total length of 2000 time steps. The network outputs were recorded at every time step, yielding predicted activity trajectories for all output neurons. To assess the dynamics to perturbations in the stimuli input, we constructed a second, ‘noisy’ input condition in which the same integer stimulus sequence was again mapped to 41-dimensional vectors, but the representation departed from strict one-hot encoding: the channel corresponding to the presented stimulus retained an activation of 1, whereas all non-stimulus channels were assigned a small constant baseline value (0.05), thereby introducing controlled, uniform signal across input channels while preserving the identity of the active stimulus. This noisy tensor, with identical shape and temporal structure, was fed through the same trained network under identical inference settings, producing a second set of predicted activity trajectories. For both conditions, the resulting predictions were measured with the same methods as before to evaluate critical temperatures and effective dimensionalities.

To quantify the effect of input noise at the single-neuron level, we computed the absolute difference between the standard and noisy predictions for each neuron and time point. To examine how input perturbations affected the emergent population structure, we next characterized the pairwise Pearson correlation between neurons predicted activity time courses separately for the standard and noisy input conditions, forming two N × N correlation matrices. From each matrix, we extracted the upper-triangular, off-diagonal entries to obtain the distribution of unique neuron–neuron correlation coefficients.

## Supplementary Materials

**Sup. Fig. 1.**
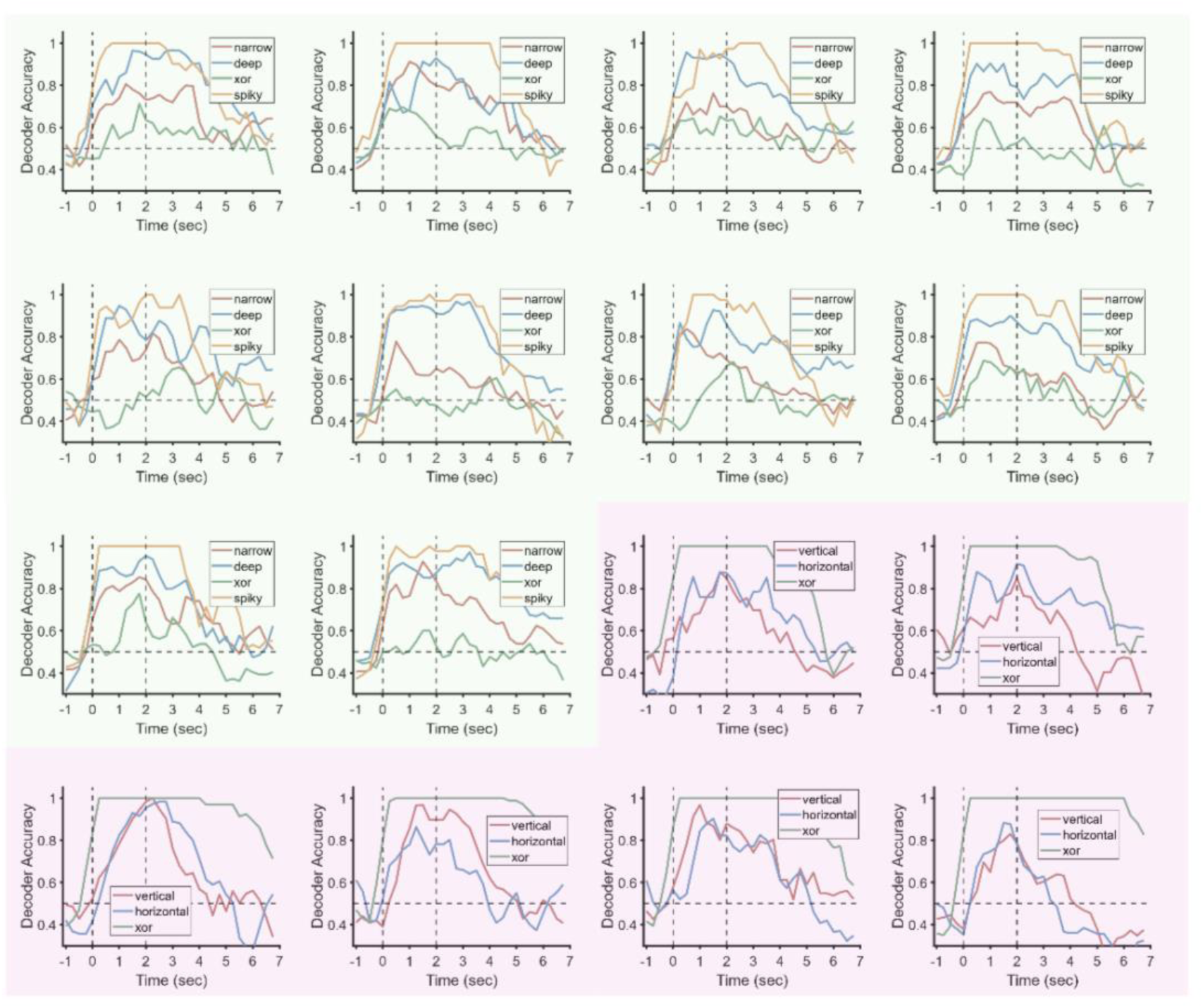
Time-resolved SVM decoding accuracy along task-defined axes in the shape and motion paradigms. Each subpanel corresponds to one experiment; background color indicates cohort membership (shape, green; motion, purple). In the shape cohort, decoding accuracies are shown for four 2×2 discriminations: narrow, deep, and XOR (classifications along the corresponding feature axes, as indicated in the legend), and spiky (diagonal category discrimination: spiky [D&N] vs. stubby [S&W]). In the motion cohort, decoding accuracies are shown for three 2×2 discriminations (horizontal, vertical, and XOR), each corresponding to classification along the indicated axis. The horizontal dashed line denotes chance accuracy (50%); vertical dashed lines mark stimulus duration. Across experiments in both cohorts, decoding accuracy increases after stimulus onset and decreases toward chance after stimulus offset.

**Sup. Fig. 2.**
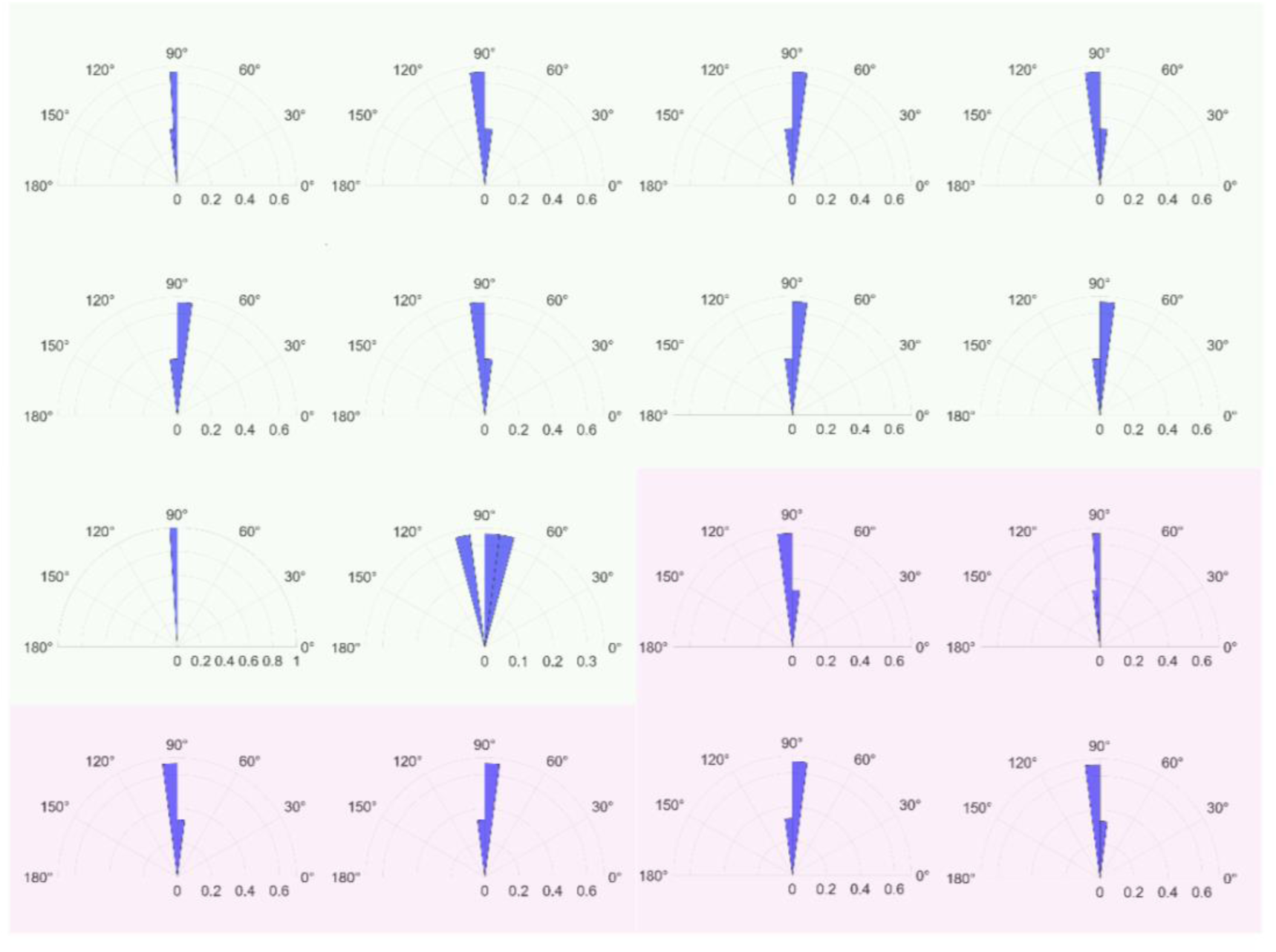
SVM axes are approximately orthogonal across repeated training runs within each experiment. Rose plots show distributions of pairwise angles between SVM axes for each experiment (N = 16). For the shape cohort (green background; N = 10), four SVM axes were obtained per training run, whereas for the motion cohort (purple background; N = 6), three SVM axes were obtained per run. Within each experiment, SVM axes were retrained 8 times independently. For each run, all pairwise angular separations among the 4 (shape) or 3 (motion) axes were computed and visualized as polar histograms. Across experiments in both cohorts, angle distributions are concentrated near 90°, indicating a stable, near-orthogonal low-dimensional axis structure across retraining.

**Sup. Fig. 3.**
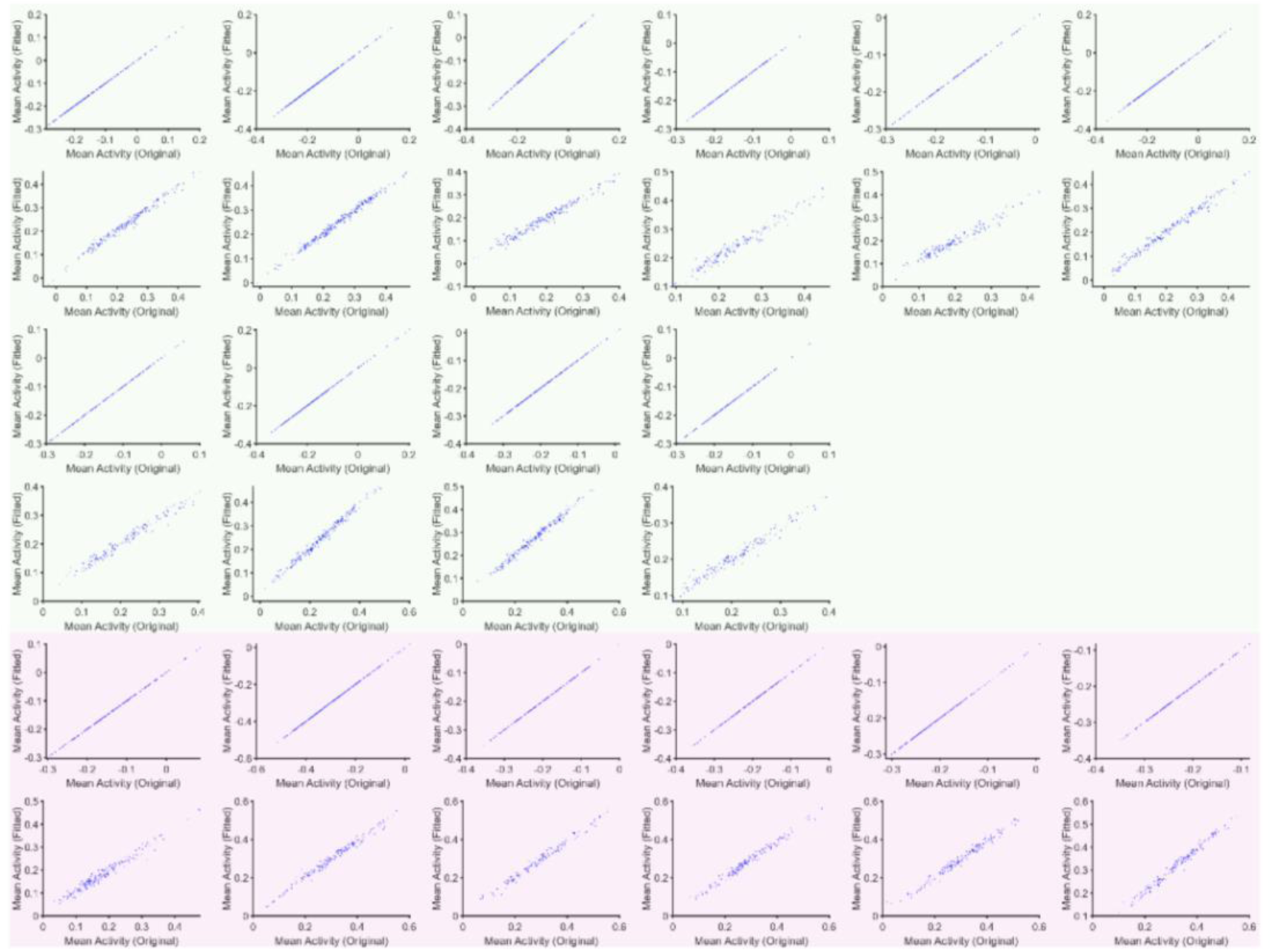
Pairwise MaxEnt models reproduce single-neuron mean activity across experiments. Model-predicted versus empirical mean activity is shown for all experiments (n = 16). In each pair of subpanels, each point corresponds to one neuron. The upper subpanel shows the stim-on epoch and the lower subpanel shows the stim-off epoch. Experiments are ordered left-to-right; background color indicates cohort membership (shape, green; motion, purple). Points lie close to the identity line (y = x), indicating that the fitted pairwise MaxEnt models capture first-order (single-neuron) statistics in both stim-on and stim-off epochs.

**Sup. Fig. 4.**
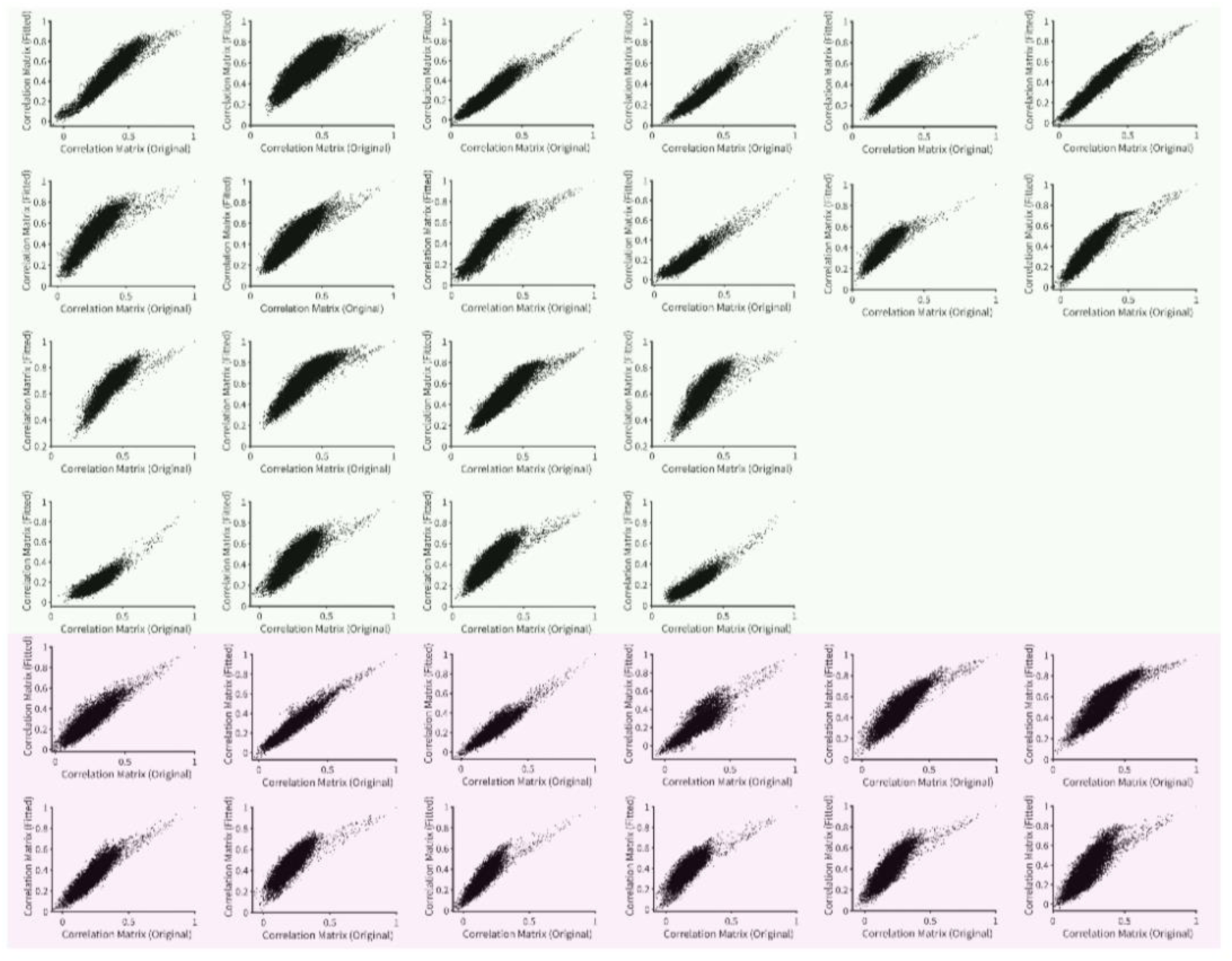
Pairwise MaxEnt models reproduce empirical pairwise correlations across experiments. Model-predicted versus empirical pairwise correlations are shown for all experiments (n = 16). In each pair of subpanels, each point corresponds to one neuron pair. The upper subpanel shows the stim-on epoch and the lower subpanel shows the stim-off epoch. Experiments are ordered left-to-right; background color indicates cohort membership (shape, green; motion, purple). Points cluster near the identity line (y = x), indicating that the fitted pairwise MaxEnt models capture second-order (pairwise) statistics in both stim-on and stim-off epochs.

**Sup. Fig. 5.**
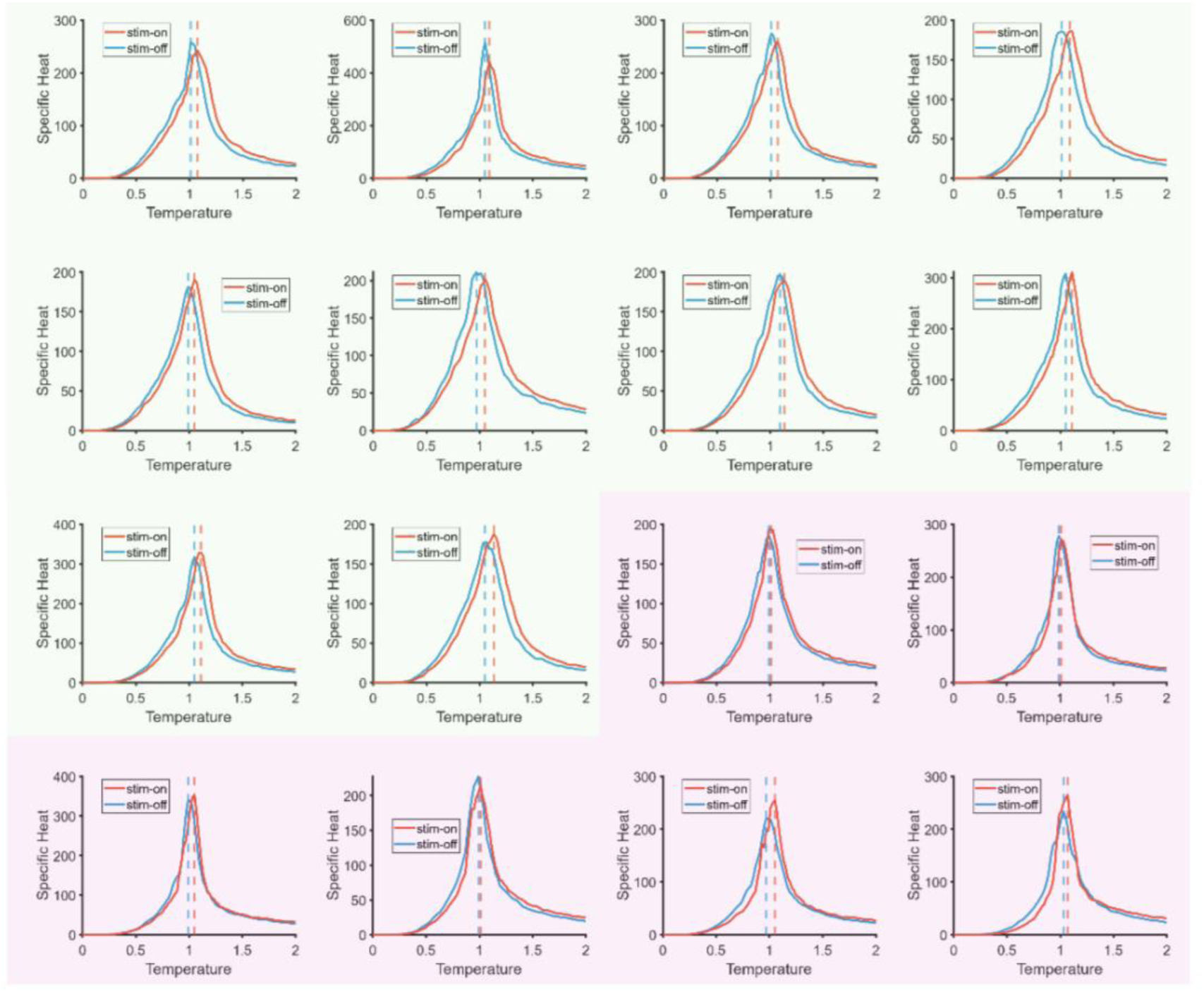
Specific heat signatures from pairwise MaxEnt models are consistent across animals in the shape and motion cohorts. Specific-heat curves were computed from pairwise maximum entropy (MaxEnt) models fitted separately to population activity during the stim-on (stimulus-evoked; red) and stim-off (spontaneous; blue) epochs, and evaluated across a range of model temperatures. Each subpanel shows one mouse; background color indicates cohort membership (shape, green; motion, purple). Dashed lines mark the specific-heat peak value(s); the corresponding temperature defines the critical temperature for that condition. Across animals in both cohorts, stim-on and stim-off activity exhibit qualitatively similar, reproducible shifts in the peak location of the specific-heat curves.

**Sup. Fig. 6.**
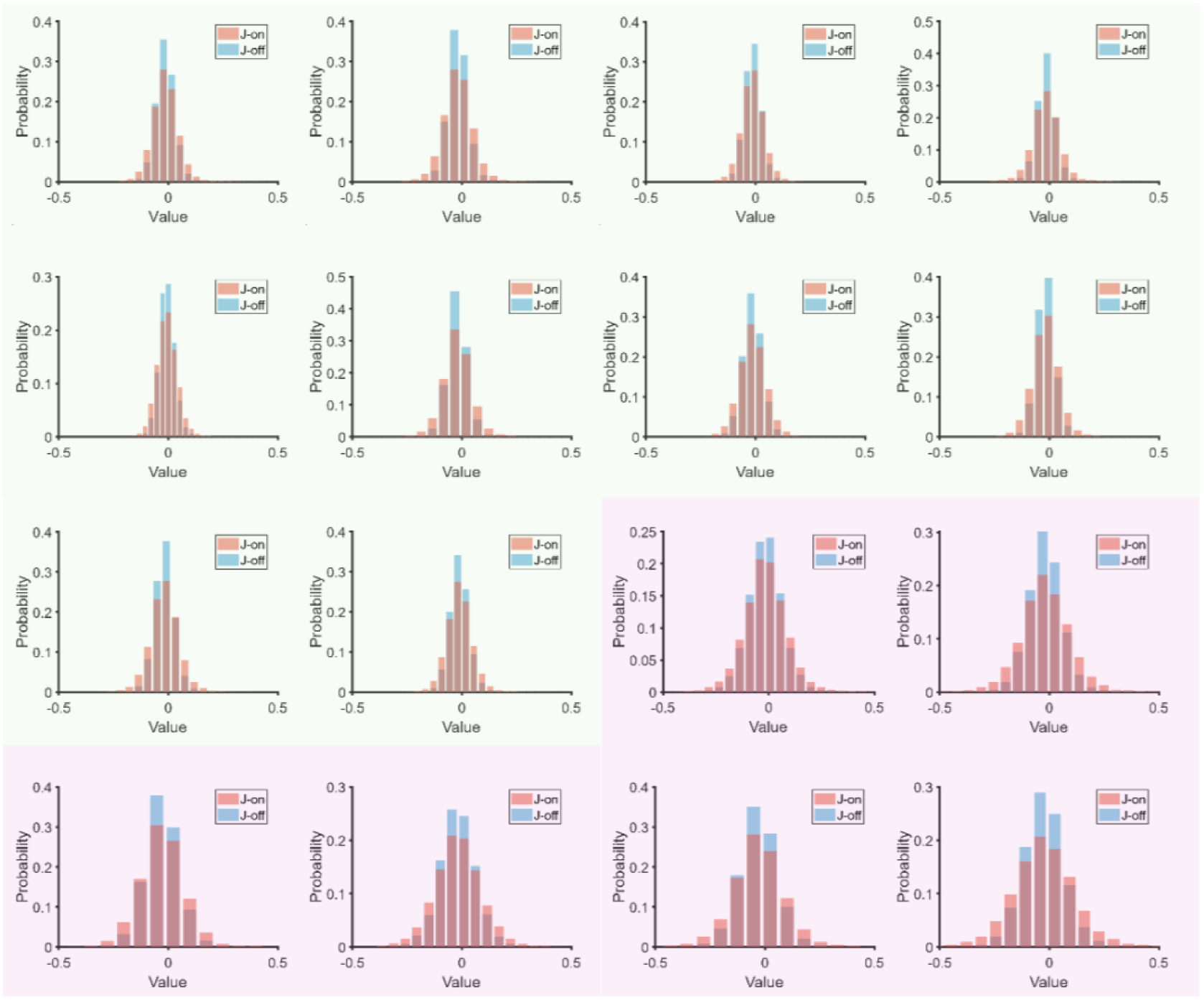
Distributions of inferred coupling parameters (J) from pairwise MaxEnt models across experiments and epochs. Probability distributions of pairwise coupling parameters J inferred from pairwise maximum entropy (MaxEnt) models fitted separately to the stim-off (spontaneous; blue) and stim-on (stimulus-evoked; red) epochs for each experiment (shape cohort: N = 10, green background; motion cohort: N = 6, purple background). Across experiments and epochs, J distributions are approximately Gaussian and centered near zero, indicating a balance of positive and negative couplings at the population level. Relative to stim-off, stim-on distributions are consistently broader, indicating an increased spread (magnitude) of inferred pairwise interactions during the stim-on epoch.

**Sup. Fig. 7.**
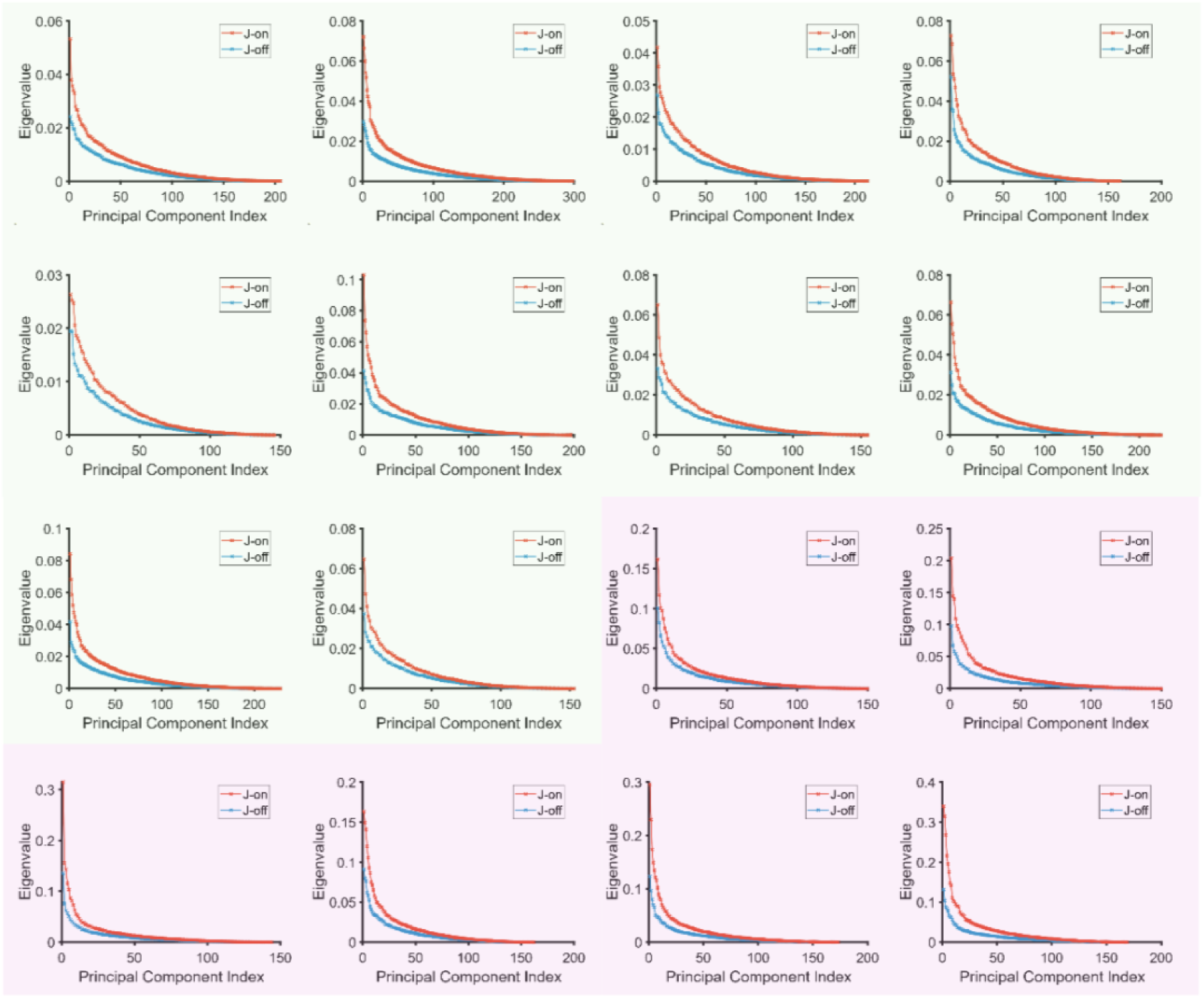
Principal-component spectra of inferred coupling matrices indicate stronger low-dimensional structure during the stim-on epoch. For each experiment, we computed the eigenspectrum of the inferred coupling matrix J (eigenvalues sorted in descending order) for stim-off and stim-on models. Each subpanel corresponds to one experiment (shape cohort, green background; motion cohort, purple background). Across experiments, the stim-on spectra (red) are systematically shifted upward relative to stim-off (blue), indicating that a larger fraction of the variance in J is captured by the leading principal components during stimulation and that eigenvalues are more concentrated in the top-ranked modes.

**Sup. Fig. 8.**
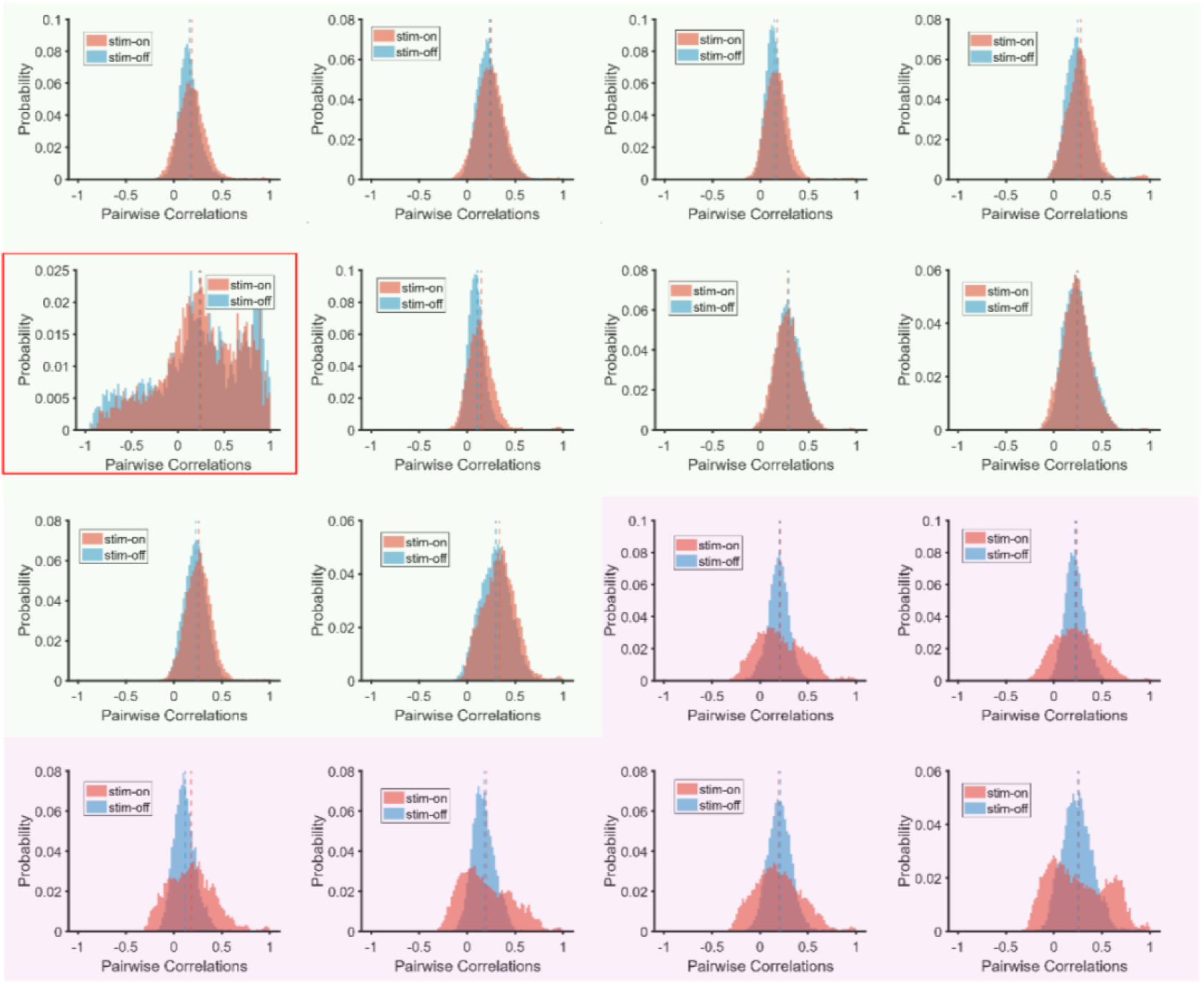
Pairwise correlation distributions across animals during stim-on and stim-off epochs. Distributions of pairwise Pearson correlations among simultaneously recorded neurons were computed separately during the stim-on (stimulus-evoked; red) and stim-off (spontaneous; blue) epochs. Each subpanel shows one mouse; background color indicates cohort membership (shape, green, N = 10; motion, purple, N = 6). Within each cohort, distribution shapes and summary statistics are reproducible across animals. One animal in the shape cohort (mouse 5; highlighted by a red box) exhibits an unusually broad distribution, attributable to a large common drift component in the recorded signals that could not be fully removed by preprocessing. Because this drift acts primarily on long timescales, it does not affect analyses performed on shorter (trial-scale) segments. For the cross-animal summary, this mouse’s distribution standard deviation was treated as an outlier and excluded from the group-level analysis of distribution width; all other analyses were performed as described.

**Sup. Fig. 9.**
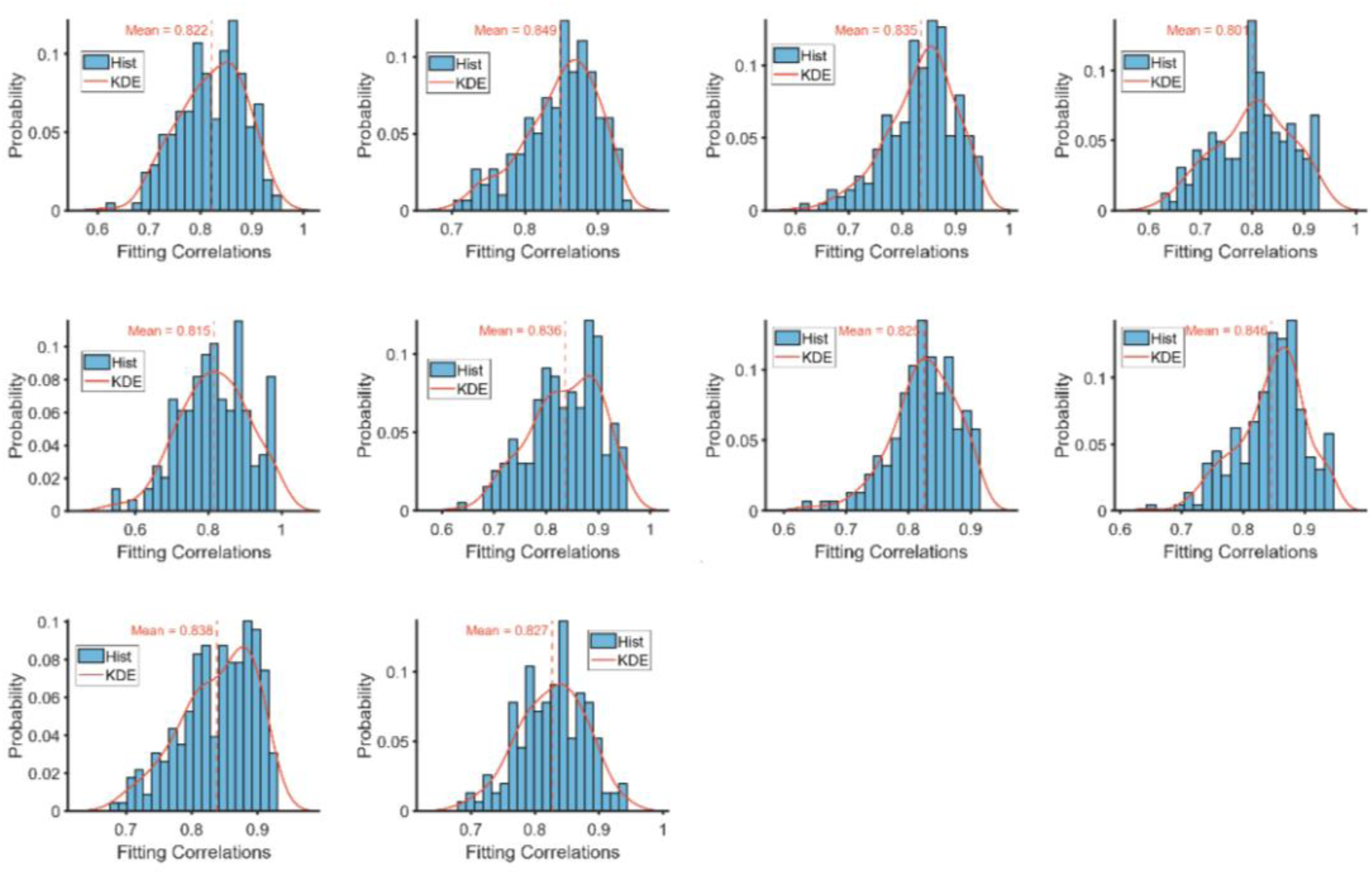
RNN predictions match single-neuron calcium dynamics across shape experiments. For each shape experiment (N = 10), Pearson correlations were computed between RNN unit activity and empirically recorded calcium traces (RNN driven by the experimental input sequence; correlations computed between predicted and observed traces). For each experiment, the distribution of correlation coefficients across all matched RNN–data unit pairs is shown as a histogram (probability distribution). Red curves indicate kernel density estimates (KDEs); vertical dashed lines denote the mean correlation for each experiment. Across experiments, mean correlations are consistently high (all > 0.8), indicating that the trained RNNs robustly reproduce single-unit temporal dynamics in the shape paradigm.

## Appendix: Relation between critical temperature and effective manifold dimension in a pairwise Ising model

For the standard pairwise maximum-entropy model, we assume no external stimulus in the Hamiltonian itself; different stimulus conditions change the inferred couplings and fields. The magnetization vector is *m* = ⟨*s*⟩. For the rest of the derivation, we focus on regimes where *m* is small and the system is in a paramagnetic phase (no spontaneous symmetry breaking).

The mean activities satisfy the usual mean-field self-consistency equation

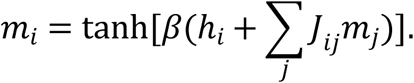

For small arguments and omit the external field, it becomes

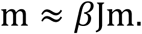

A non-zero solution appears when

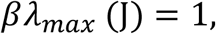

Defining the critical inverse temperature β_*c*_ = 1/T_c_

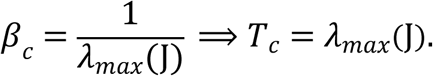

So, the critical temperature is equal to the largest eigenvalue of the coupling matrix. Any increase in the top eigenvalues of J (e.g. under stim-on) raises *T_c_*.

The susceptibility gives the standard linear-response relation

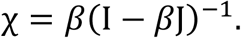

For an Ising model, fluctuation–response yields

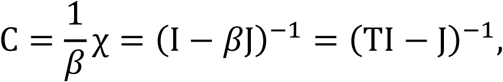

so, the covariance eigenvalues are

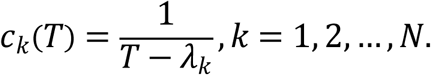

In the eigenbasis of J, the covariance eigenvalues satisfy *c*_*k*_(*T*) ∝ 1/(*T* − λ_*k*_) (equivalently *c*_*k*_(*T*) = *T*/(*T* − λ_*k*_) up to a T-dependent prefactor that is common across modes). As T approaches the largest eigenvalue λmax, the leading covariance mode diverges, reducing the effective dimensionality defined by the participation ratio. Strictly speaking, below *T_c_* the stable mean activities satisfy *m*^∗^ ≠ 0, and the linear response around this state involves a diagonal gain factor *D* = *diag*(1 − *m*^∗2^), yielding *C* = [*I* − *T*^−1^*DJ*]^−1^*D*. For simplicity, and we assume magnetizations are small, *D* ≈ *I*, which reduces the covariance to *C* ≈ (*TI* − *J*)^−1^. All results on the relation between *T_c_*, the spectrum of *J*, and the effective dimension *D*_*eff*_ are to be understood in this weakly magnetized mean-field regime.

For a fixed stimulus condition, the Ising model defines a cloud of neural activity vectors σ. Around its mean *m*, this cloud is locally well approximated by a Gaussian ellipsoid with covariance *C*. Let *R*_*k*_ be the principal radii of this local ellipsoid (square of the radius along eigenmode k is proportional to *c*_*k*_). In Chung et al.’s manifold anchor geometry, the Gaussian manifold dimension of such an ellipsoid is defined as a participation-ratio measure of the radii:

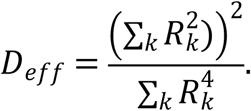

If we take

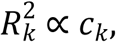

the proportionality cancels between numerator and denominator, and the effective dimension becomes

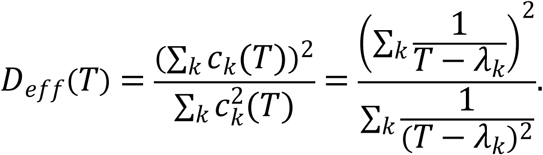

assuming ordering λ_1_ ≥ λ_2_ ≥ ⋯ ≥ λ_*N*_, and write the spectrum in units of *T_c_*:

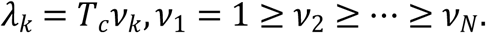

Then the covariance eigenvalues at temperature T are

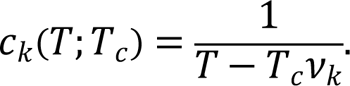

Define

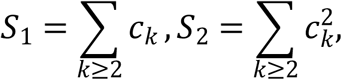

Then the effective dimension can be written as

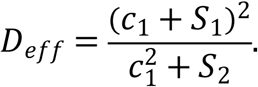

Treat *S*_1_ and *S*_2_ as slowly varying with respect to *c*_1_. Differentiate *D*_*eff*_ with respect to *c*_1_:

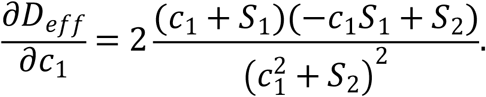

When the leading variance dominates the others, we have −*c*_1_*S*_1_ + *S*_2_ < 0, hence

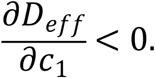

At the same time,

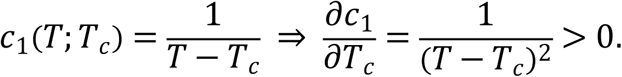

Thus,

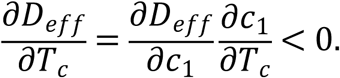

Even under the finite-size/weak-magnetization approximation, it can still serve as a heuristic explanation of the spectrum–dimension relationship.

